# Emergence of a novel immune-evasion strategy from an ancestral protein fold in bacteriophage Mu

**DOI:** 10.1101/861617

**Authors:** Shweta Karambelkar, Shubha Udupa, Vykuntham Naga Gowthami, Sharmila Giliyaru Ramachandra, Ganduri Swapna, Valakunja Nagaraja

## Abstract

The broad host range bacteriophage Mu employs a novel ‘methylcarbamoyl’ modification to protect its DNA from diverse host restriction systems. Biosynthesis of the unusual modification is a longstanding mystery. Moreover, isolation of Mom, the phage protein involved in the modification has remained elusive to date. Here, we characterized the co-factor and metal binding properties of Mom and provide a molecular mechanism to explain ‘methylcarbamoyl’ation by Mom. Our computational analyses revealed a conserved GNAT (GCN5-related N-acetyltransferase) fold in Mom, predicting acetyl CoA as its co-factor. We demonstrate that Mom binds to acetyl CoA and identify the active site. Puzzlingly, none of the > 309,000 GNAT members identified so far catalyze Mom-like modification of their respective substrates. Besides, conventional acid-base catalysis deployed by typical acetyltransferases cannot support methylcarbamoylation of adenine seen in Mu phage. In contrast, free radical-chemistry, catalyzed by Fe-S cluster or transition metal ions can explain the seemingly challenging reaction between acetyl CoA and DNA. We discovered that Mom is an iron-binding protein, with the Fe^2+/3+^ ion colocalized with acetyl CoA in the active site of Mom. Mutants defective for binding Fe^2+/3+^ or acetyl CoA demonstrated compromised activity, indicating their importance in the DNA modification reaction. Iron-binding in the GNAT active site is unprecedented and represents a small step in the evolution of Mom from the ancestral acetyltransferase fold. Yet, the tiny step allows a giant chemical leap from usual acetylation to a novel methylcarbamoylation function, while conserving the overall protein architecture.

**Summary:** Studying the arms race between bacteria and their viruses (bacteriophages or phages) is key to understanding microbial life and its complexity. An unprecedented DNA modification shields phage Mu from bacterial restriction endonucleases that destroy incoming phage DNA. Nothing is known of how the modification is brought about, except that a phage protein Mom is involved. Here, we discover acetyl CoA and iron as key requirements for the modification. We explain how by evolving the ability to bind iron - a transition metal capable of generating highly reactive free radicals, a well-studied scaffold like the acetyltransferase fold can gain novel catalytic prowess in Mom. These findings have broad implications for gene editing technologies and therapeutic application of phages.

## Introduction

In the ever-escalating arms race against bacteria, phages appear to be edging out their hosts by engineering a number of strategies ^1,2^. For example, phage Mu has evolved an immune-evasion strategy in the form of a unique and broadly protective DNA modification. The methylcarbamoyladenine modification or ncm^6^A, confers resistance to >20 diverse restriction enzymes, enabling Mu to invade multiple bacterial hosts in addition to *Escherichia coli* ^3-7^ (Figure 1A). ncm^6^A is unique to Mu, being found in no other viral or cellular life form. Decades after its discovery, little is known about the modification. Overexpression of Mom, the phage protein catalyzing the DNA modification, is lethal to *E. coli* ^8,9^. The toxicity also explains why phage Mu limits *mom* expression to a brief period in late lytic infection, after phage DNA replication is completed and the host is destined to die ^10^. DNA modification by Mom enables dramatically higher (up to 10^4^ fold) efficiencies of plating compared to Mu *mom*^-^ phages on restricting strains of *E. coli* ^4^. Owing to the selective advantage, numerous host and phage-encoded proteins have evolved to regulate the potentially lethal *mom* gene in diverse ways ^4,10,11^. While intricate details surrounding *mom* gene regulation have been teased apart, the biochemistry of Mom and molecular mechanisms underlying the modification are still shrouded in mystery. Moreover, the absence of structurally or functionally characterized homologs of Mom has precluded confident bioinformatic prediction of the enzymatic requirements of Mom and the chemistry underlying the biosynthesis of this unusual, bulky DNA modification ^12^. Here, we present the first successful isolation and characterization of Mom, including the discovery of its active site, metal-binding properties and co-factor requirements. Based on these findings, we explain how phage Mu may have stumbled upon a novel recipe for immune-evasion using ancient molecular ingredients.

**Figure 1.**
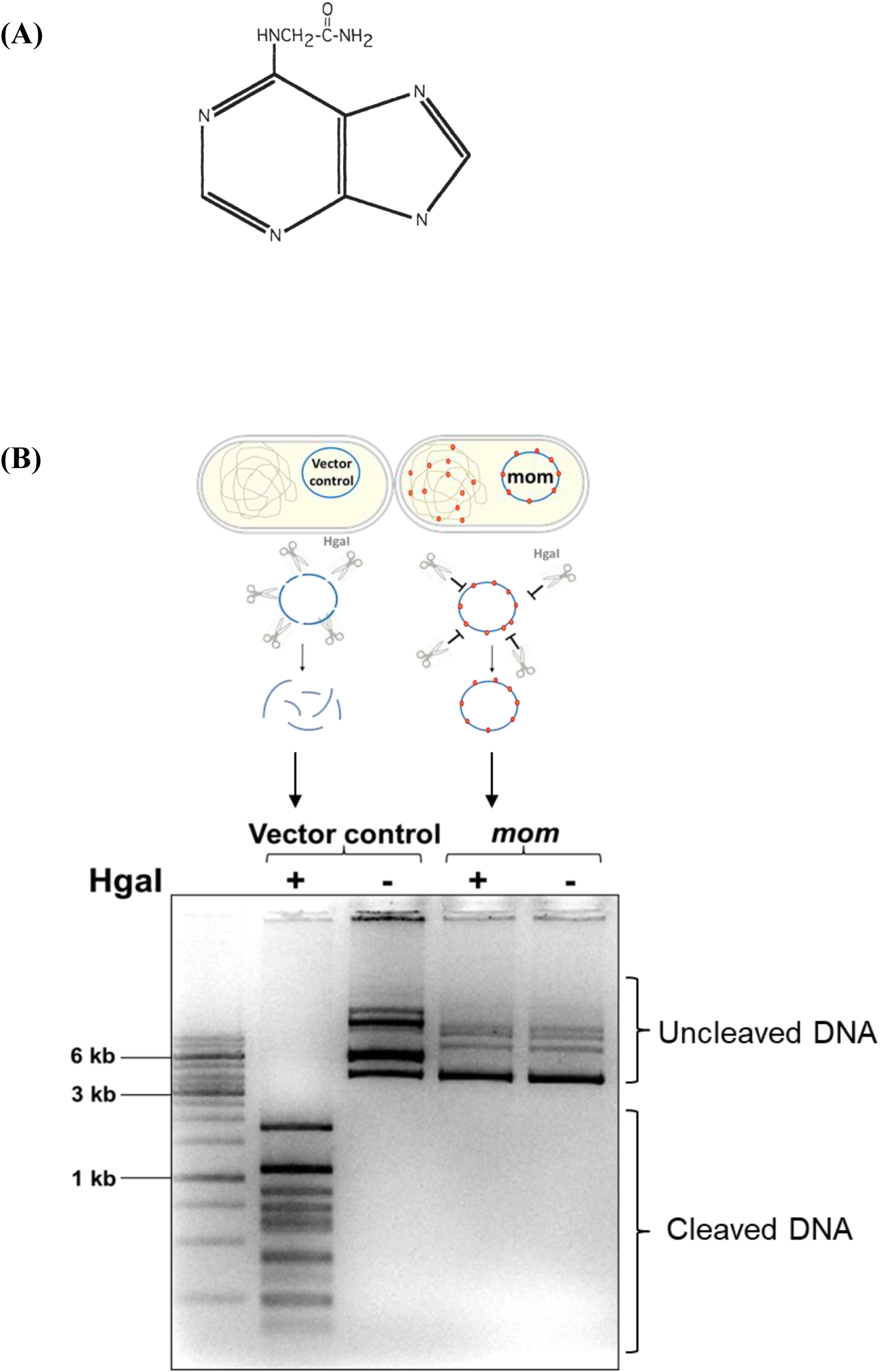
DNA modification activity of Mom. (A) Structure of the modified base *N*^6^-methylcarbamoyl adenine. (B) Analysis of the *in vivo* DNA modification activity of Mom using HgaI. Plasmid DNA was isolated from *E. coli* C41 cells harbouring the empty vector (vector control) or expressing Mom as depicted in the schematic. To examine if cellular DNA was modified by Mom, 1 μg plasmid DNA was digested with HgaI and products were analyzed on a 1% agarose ethidium bromide gel. While plasmids from vector control cells get digested into expected sized products, plasmids from *mom* expressing cells remain unaffected by HgaI.

## Results

### Expression and activity assay for Mom

Isolation of Mom remained elusive in the past owing to protein toxicity ^8^ and insolubility upon overexpression in *E. coli* ^9^. To successfully clone and express Mom in a soluble form, we deployed a tightly regulated, dual plasmid pET expression system developed previously ^13^. Mom-modified DNA from phage Mu-infected hosts is known to be resistant to at least 20 restriction endonucleases, including HgaI, PvuII, and SalI, to which unmodified DNA is sensitive ^6^. Taking cue from this, we developed a convenient and robust assay for detecting Mom activity. To determine whether Mom, overexpressed using the pET system, was proficient at modifying DNA *in vivo*, we isolated plasmid DNA from cells expressing Mom and treated it with the restriction endonuclease HgaI. While DNA from vector-control cells was completely digested to the expected cleavage products, DNA from cells that expressed Mom remained undigested, indicating that cloned Mom is functional *in vivo* (Figure 1B). Similar results were obtained with PvuII and SalI digestion of plasmid DNA isolated from Mom-expressing cells (Supplementary Figure S1). Furthermore, single point mutations in Mom (described in later sections) abolish the resistance to HgaI, indicating that protection from HgaI is specific to Mom activity.

### Properties of Mom

Having confirmed the activity of Mom *in vivo*, we next attempted to isolate Mom. Hexahistidine-tagged Mom purified by Ni-NTA chromatography precipitated upon removal of imidazole. Hence, we designed an alternate ion-exchange-based protocol for purifying Mom in a soluble form (Supplementary Figures S2A and S2B). The molecular mass of untagged Mom, analyzed by LC-ESI-MS, matched the expected mass of 27,050 Da and revealed that the N-terminal methionine encoded by the gene is absent (Supplementary Figure S2C). The far-UV CD spectrum of Mom showed that it has appreciable secondary structure comprised of alpha helices and beta sheets (Supplementary Figure S2D). The fluorescence spectrum of the protein contained an expected red shift and a change in emission intensity upon chemical denaturation (Supplementary Figure S2E), indicating that in the native state the protein is folded with some buried tryptophan residues. Surface plasmon resonance studies showed that Mom binds to DNA with an affinity of ∼0.2 nM (Supplementary Table S1). In addition, analytical gel filtration and glutaraldehyde cross-linking experiments indicated that Mom exists as a dimer in solution (Supplementary Figures S2F and G). Having successfully expressed functionally active Mom and characterized its properties, we set out to understand its enigmatic enzyme function.

### Acetyltransferase fold in Mom

Given that no structurally or functionally characterized homologs of Mom exist, we subjected the sequence of Mom to *de novo* and threading-based structure-prediction tools Robetta and I-TASSER, respectively. We found an evolutionary relationship between Mom and the GCN5-related N-acetyltransferase (GNAT) superfamily, as reported previously ^12^ (Figure 2). The analyses predict a classic GNAT fold occupying the core of Mom, with extensions at the N- and C-termini (Figure 2). The GNAT fold of Mom is comprised of a central, highly curved beta sheet sandwiched between five alpha helices. In addition to the overall fold, Mom displays several GNAT-like structural features, including the classic splayed beta strands β4 and β5 and a resultant V-shaped cleft for binding of acyl coenzyme A ^14^ (Figure 2). Given the structural similarity with acetyltransferases, the COFACTOR server predicted acetyl CoA, the co-factor most commonly utilized by the GNAT superfamily, as the co-factor of Mom (Figure 2). As in prototypical acetyltransferases, acetyl CoA in the Mom-acetyl CoA model fitted into the structurally conserved V-shaped cleft of Mom, adopting a characteristic ‘bent’ conformation (Figure 2), again suggesting acetyl CoA as the co-factor of Mom ^15^.

**Figure 2.**
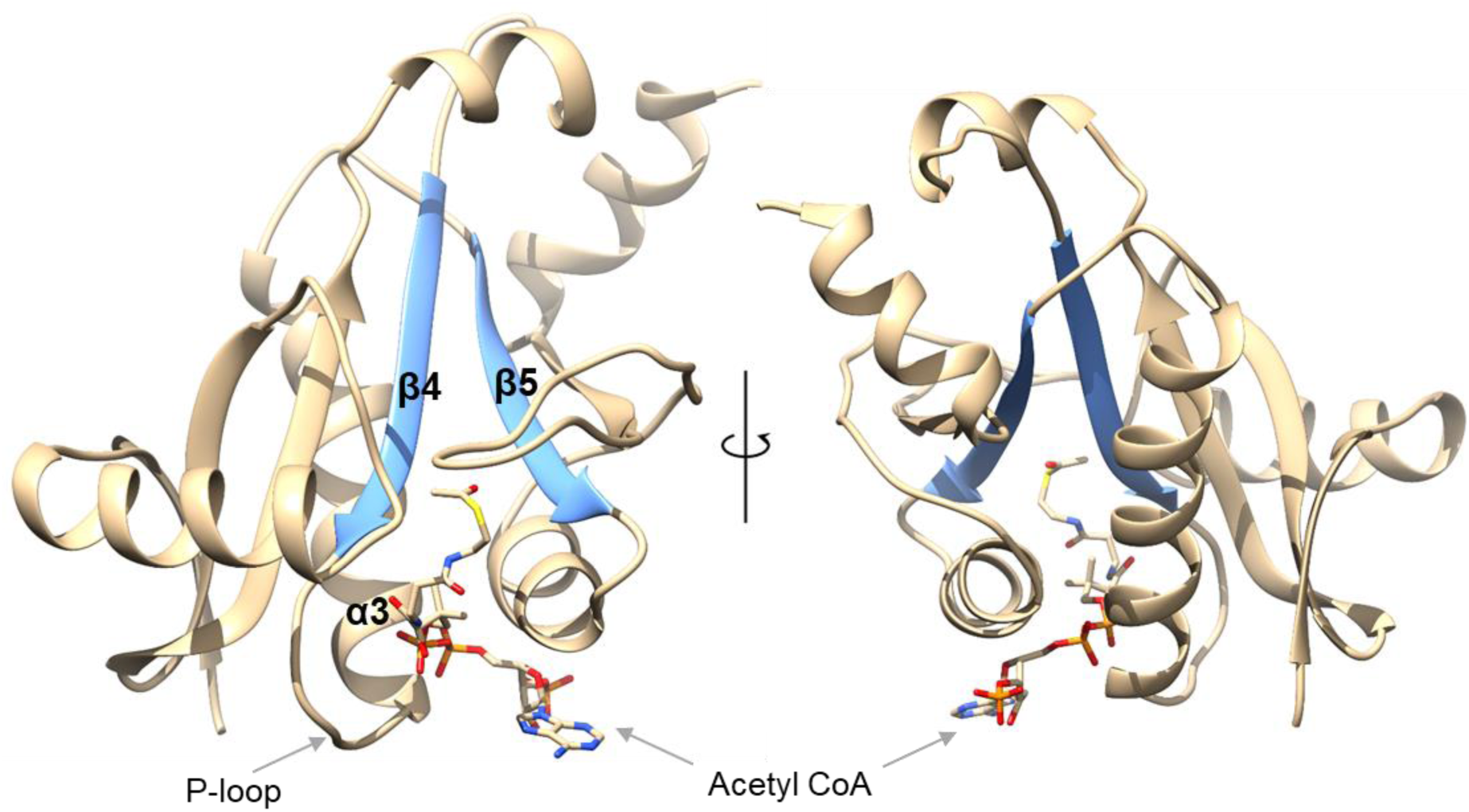
Mom is structurally related to acetyltransferases. Mom harbors a classic GNAT fold in its core, as revealed by disparate structure prediction methods. The fold comprises a six-stranded curved beta sheet sandwiched between five alpha helices. The beta sheet is largely antiparallel; the only exception being strands β4 and β5 (highlighted in blue), a structural feature conserved in the GNAT superfamily. β4 and β5 splay apart towards their C-termini, forming an inverted V-shaped cleft (blue), likely occupied by acetyl CoA, the co-factor most commonly employed by the GNAT superfamily, bound in a characteristic bent conformation. In addition, Mom harbors the P-loop and the βαβ (β4-α3-β5) motif, structural elements critical for generating the acetyl CoA binding site in the GNAT fold, as indicated ^14^. As in prototypical GNAT structures, the pyrophosphate binding loop or P-loop of Mom appears poised to form direct or water-mediated hydrogen bonds to the pyrophosphate arm of acetyl CoA, while the backbone of β4 interacts intimately with the pantothenate arm of acetyl CoA via hydrogen bonds.

### Mom binds to acetyl CoA

To test if Mom interacts with acetyl CoA, we employed differential scanning fluorimetry, which detects changes in the melting temperature (Tm) of a target protein upon interaction with a ligand ^16,17^. Interaction between Mom and acetyl CoA was evident from the dose-dependent effect of acetyl CoA on the Tm of Mom. (Figure 3A and Supplementary Table S2). In contrast, no such effect was observed with compounds like S-adenosyl methionine (SAM), NAD^+^, glycine, ATP, etc. (Figure 3B and Supplementary Table S2). A decrease in the Tm of Mom upon binding to acetyl CoA indicated that the interaction results in a more flexible or open protein conformation. A comparable effect was also observed with coenzyme A, a by-product of acyl CoA hydrolysis. Longer CoA derivatives exhibited weaker or no interaction with Mom (Supplementary Table S2). Such a preference for acetyl CoA over longer coenzyme A derivatives is displayed by other acetyl CoA-utilizing GNAT members as well ^18^. We next quantified the Mom-acetyl CoA interaction using microscale thermophoresis, which measures biomolecular interactions by monitoring the motion of molecules along microscopic temperature gradients, detecting ligand-induced changes in the hydration shell, charge, or size of macromolecules ^19,20^. While Mom bound to acetyl CoA with K_d_ ∼ 49 μM, no detectable interaction was observed with the control ligand S-adenosyl methionine (Figure 3C).

**Figure 3.**
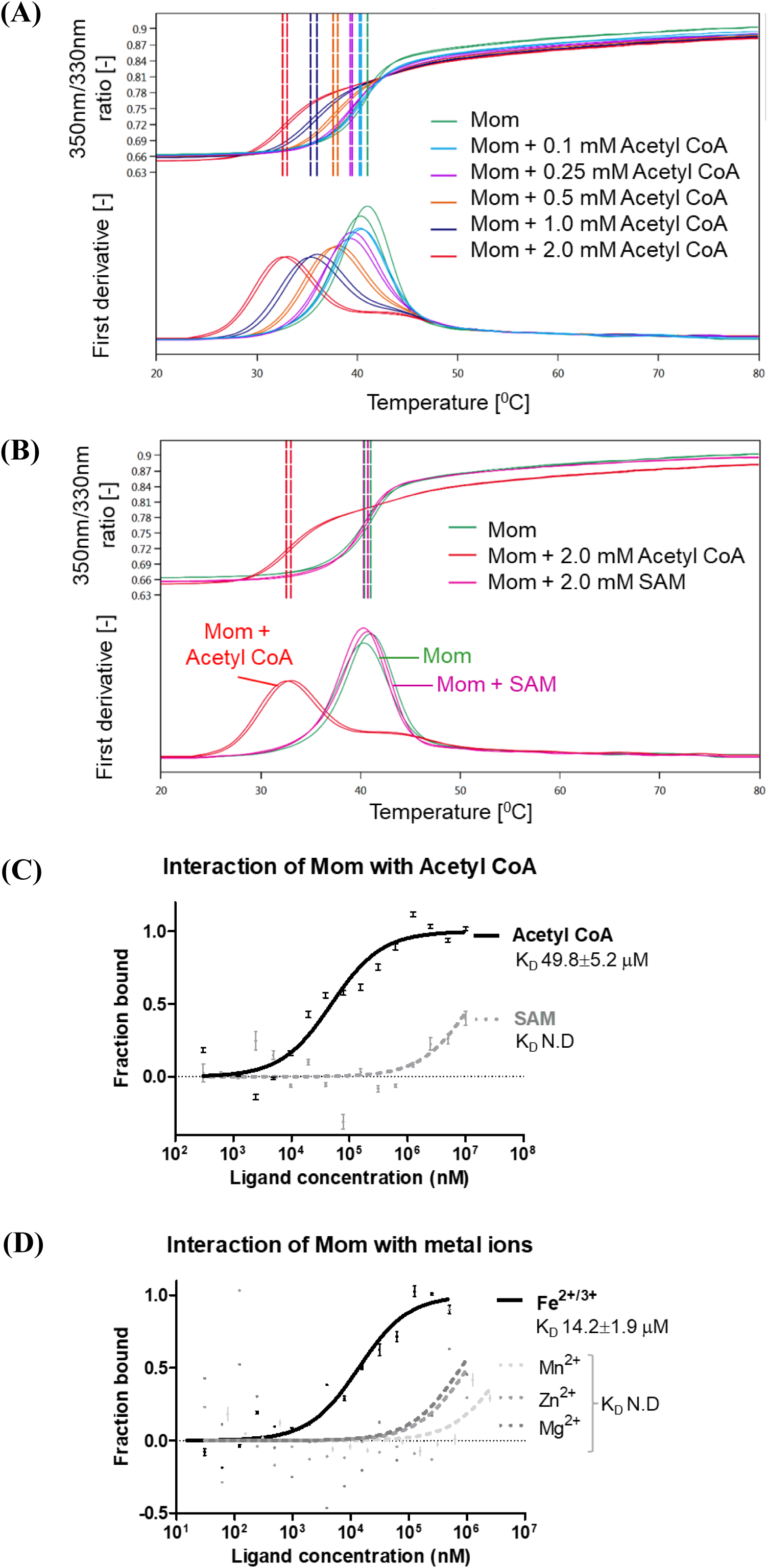
Interaction of Mom with acetyl CoA and other ligands. (A) Dose-dependent effect of acetyl CoA on the thermostability of Mom. Mom was incubated with indicated concentrations of acetyl CoA and subjected to thermal ramp (1°C/min, 20-80°C). The shift in intrinsic tryptophan fluorescence of proteins upon temperature-induced unfolding was monitored by detecting the emission fluorescence at 330 and 350 nm. Protein melting points (Tm) were determined from the transition midpoints (dotted vertical lines in the upper panel) corresponding to peaks in the first derivatives of the 350/330 fluorescence data (lower panel). Reactions were carried out in duplicates. **(B) Comparison of thermostability of Mom in the presence of acetyl CoA and SAM.** Representative thermal denaturation curves of Mom alone (green), Mom incubated with 2 mM acetyl CoA (red) and Mom incubated with 2 mM SAM (magenta). While SAM has no detectable effect on the thermostability of Mom, acetyl CoA significantly destabilizes Mom. Reactions were carried out in duplicates. **(C) Quantification of the binding affinity of Mom with acetyl coenzyme A using microscale thermophoresis.** Mom was fluorescently labeled and titrated against indicated concentrations of ligands. Normalized fluorescence is plotted for analysis of thermophoresis. Data were fitted using the law of mass action to determine dissociation constants (K_D_). Data shown are representative of three independent experiments. Error bars represent standard deviations of n=3 measurements. Acetyl CoA interacts with Mom while SAM shows undetectable interaction. **(D) Specificity of Mom-iron interaction.** Microscale thermophoresis was carried out using fluorescently labeled Mom and various divalent metal ions and data plotted as described for Figure 3C. While Fe^2+/3+^ interacts with Mom, Mn^2+^, Zn^2+^ and Mg^2+^ showed undetectable interaction. Metal ions Ca^2+^, Co^2+^ and Cu^2+^ also failed to interact with Mom (data could not be fitted).

### Mom catalyzes methylcarbamoyl transfer, not acetyl transfer

Based on the structural resemblance of Mom to acetyltransferases and its ability to bind acetyl CoA, it was tempting to hypothesize that Mom functioned as a prototypical acetyltransferase. However, Mom is known to transfer an unusual methylcarbamoyl (-CH_2_CONH_2_) group rather than acyl group (-COR, such as acetyl (-COCH_3_), propionyl (-COC_2_H_5_), etc.) typically transferred by acetyltransferases ^3,14^. To confirm that the modification was indeed ncm^6^A and not some acylated form of adenine, we revisited the original study by Swinton et al. ^3^. To do so, genomic DNA from *E. coli* cells expressing Mom was digested to nucleosides and analyzed using LC-ESI/MS^n^. In the *mom*^+^ samples, we observed a unique species (MomdA), with mass corresponding to ncm^6^dA (methylcarbamoyldeoxyadenosine), accompanied by reduced levels of unmodified deoxyadenosine (dA) (Supplementary Figures S3A and B). Furthermore, collision-induced dissociation of MomdA yielded fragments diagnostic of ncm^6^dA (Supplementary Figures S3C and D), thus reaffirming the identity of the modification. The fragmentation data ruled out acetyl(-COCH_3_)adenine and glycyl(-COCH_2_NH_2_)adenine (identical mass as ncm^6^A), conceivable with known GNAT chemistries employing acetyl CoA and glycyl tRNA, respectively ^14,21^. Having confirmed the chemical nature of the modification, we next sought to understand how Mom could accomplish the methylcarbamoyl transfer.

### Exploring conventional and unconventional chemistries to decipher Mom mechanism

Acetyl CoA participates as a two-carbon donor by donating its acetyl group, engaging via the acetyl moiety’s carbonyl carbon (e.g. GNAT mechanisms) or methyl carbon (e.g. condensation reactions) ^15,22^. As stated before, GNAT mechanisms support transfer of acetyl moiety (-COCH_3_) to adenine, but not of –CH_2_COR (R being -OH or -NH_2_) required for ncm^6^A ^15^. On the other hand, condensation reactions can transfer –CH_2_COR group to electrophilic acceptors but not to nucleophilic acceptors like the N^6^ amino group of adenine ^22^. This is because condensation involves deprotonation of the methyl group of acetyl CoA, which renders the carbon nucleophilic and thus non-reactive with the nucleophilic N^6^ amino group of adenine (Supplementary Figure S4A)^22^. In other words, no conventional mechanism supports acetyl CoA as the co-factor of Mom. So, we broadened our investigation and examined other potential co-factors and mechanisms for methylcarbamoylation of adenine.

Given that our *in vivo* assays demonstrated that methylcarbamoylation of DNA *in vivo* required no phage gene apart from Mom (Figure 1B), it was reasonable to hypothesize that the co-factor used by Mom must be a host-derived metabolite. We searched the *E. coli* metabolome database for molecules wherein a good leaving group is attached to the methylcarbamoyl moiety, thus favoring a direct nucleophilic attack from the substrate amine of adenine ^23^. However, no such donor candidate could be found, making the possibility of a single-step reaction unlikely. Next, we examined the involvement of potential two-carbon donors, such as glyoxylate and cx-SAM, a carboxylated derivative of S-adenosyl-L-methionine which donates methylcarboxyl group (-CH_2_COOH) to hydroxyuridine in tRNA modification reactions ^24,25^. However, Mom activity remained unaffected in strains lacking glyoxylate or the cx-SAM synthetic machinery (Supplementary Figure S4B), thus ruling out their participation in the Mom reaction.

While ncm^6^A is unique to phage Mu DNA, a similar modification ncm^5^U is common in archaeal and eukaryotic tRNA ^26^. Strikingly, Elp3 - the enzyme catalyzing the ncm^5^U modification, also harbors an acetyl CoA-binding GNAT domain ^27,28^. Additionally, Elp3 contains a radical SAM (rSAM) domain harbouring a 4Fe-4S cluster and S-adenosyl methionine (SAM) ^28,29^. A highly reactive ‘acetyl CoA radical’ produced by the rSAM domain attacks the substrate uridine, forming an adduct whose hydrolysis and amidation yields ncm^5^U ^26,28^. Although Mom lacks a rSAM domain, the presence of an acetyl CoA-binding GNAT domain hinted at an Elp3-like free radical-based mechanism for Mom. Given that some rSAM enzymes utilize trans-encoded rSAM domains ^30,31^, we hypothesized that Mom recruited a host-encoded rSAM domain. We tested this possibility by checking the ability of Mom to modify DNA in various *E. coli* strains lacking rSAM genes. An exhaustive search of the *E. coli* genome yielded 67 experimentally validated or predicted 4Fe-4S cluster genes typical of rSAM proteins. Surprisingly, Mom activity was detected in all 67 rSAM single gene knock-outs, indicating that none was necessary for Mom function and rendering them unlikely partners of Mom (Supplementary Figure S4C and Supplementary Table S3).

### Investigating genetic interactions of *mom* with the host

The inability of candidate-based approaches to identify host genes or co-factors involved in Mom activity prompted us to undertake an unbiased genome-wide approach. Multi-step reactions, often catalyzed by multi-protein complexes, alleviate the need for ‘one-step donors’ of bulky chemical moieties ^24,27,32-34^. Hence, we examined using both genetic and biochemical approaches, the possibility of Mom interacting with host protein(s). The genetic approach entailed expression of the toxic Mom protein in random transposon mutagenesis libraries of *E. coli*, followed by selection for survivors (Supplementary Figures S4D and E). Survival, despite Mom overexpression, would indicate compromised Mom function, possibly due to disruptive insertions in genes encoding the co-factor synthesis machinery or putative interacting partner(s) of Mom. However, the survivors we obtained from the assay either bore deletions in the *mom* gene or seemed to retain *mom* and the DNA modification while overcoming toxicity in unknown ways. Inability to obtain bacterial survivors that were *mom*^+^ and yet defective for DNA modification after multiple attempts suggested essentiality of or redundancy in the genes participating in the Mom modification pathway.

### Host protein interactions reveal Mom as an iron-binding protein

We next undertook a more direct biochemical approach to reveal host factors required for the modification. In a protein-protein interaction assay to capture interacting partners of Mom from *E. coli* cell extracts, we recovered ferritin and subunits of the pyruvate dehydrogenase (PDH) complex. Ferritin is an iron storage protein of *E. coli* involved in supplying iron atoms to iron-binding proteins and Fe-S cluster biosynthesis proteins ^35,36^, whereas the PDH complex is central to acetyl CoA biosynthesis ^37^. These results not only returned our attention to acetyl CoA, but also revived the possibility of a radical-based mechanism for the participation of acetyl CoA in the Mom reaction. Fe^2+^ and other transition metal ions (Mn^2+^,Cu^2+^, Co^2+^, etc.) in many metalloproteins generate highly reactive substrate radicals ^38,39^. The association of Mom with the iron-supplying protein, ferritin, as suggested by the pull-down assay, prompted us to check if Mom was an iron-binding protein. Indeed, the colorimetric ferrozine assay revealed intrinsically bound Fe^2+/3+^ in Mom (Supplementary Figure S5). Furthermore, microscale thermophoresis experiments showed that Mom binds specifically to iron and to no other divalent metal ions tested, including Mn^2+^, Cu^2+^, Mg^2+^, Co^2+^, Zn^2+^ and Ca^2+^ (Figure 3D). In retrospect, the presence of intrinsically bound iron in Mom was also indicated by the pale brown color of the purified Mom protein.

Given the well-known role of Fe^2+/3+^ in generating free radicals and accomplishing seemingly prohibitive reactions, we hypothesized a plausible scheme of reactions for Mom (Figure 4) ^40,41^. Briefly, oxidation of acetyl CoA catalyzed by Fe^2+^ generates glyoxylic acid ester of coenzyme A, which forms a Schiff base with N^6^ amine of adenine. Reduction of the Schiff base transfers the activated carboxymethyl (-CH_2_COSCoA) group to adenine, amidation of which yields ncm^6^A (Figure 4). Lack of knowledge of additional host factors or special conditions required, if any, have precluded *in vitro* reconstitution of the Mom reaction. Supplementing the reactions with alpha-ketoglutarate and ascorbate, commonly employed as reductants by Fe^2+/3+^-utilizing dioxygenases or adding whole or fractionated cell extracts failed to show any effect ^42^. Nevertheless, having observed specific interactions of Mom with acetyl CoA and Fe^2+/3+^ *in vitro*, we hypothesized that mechanisms involving acetyl CoA as the two-carbon donor and iron as a catalyst require the two reactants to be in physical proximity within the active site of Mom. We thus sought to characterize the active site of Mom by identifying and disrupting the binding sites of acetyl CoA and Fe^2+/3+^ and examining the effects on Mom activity.

**Figure 4.**
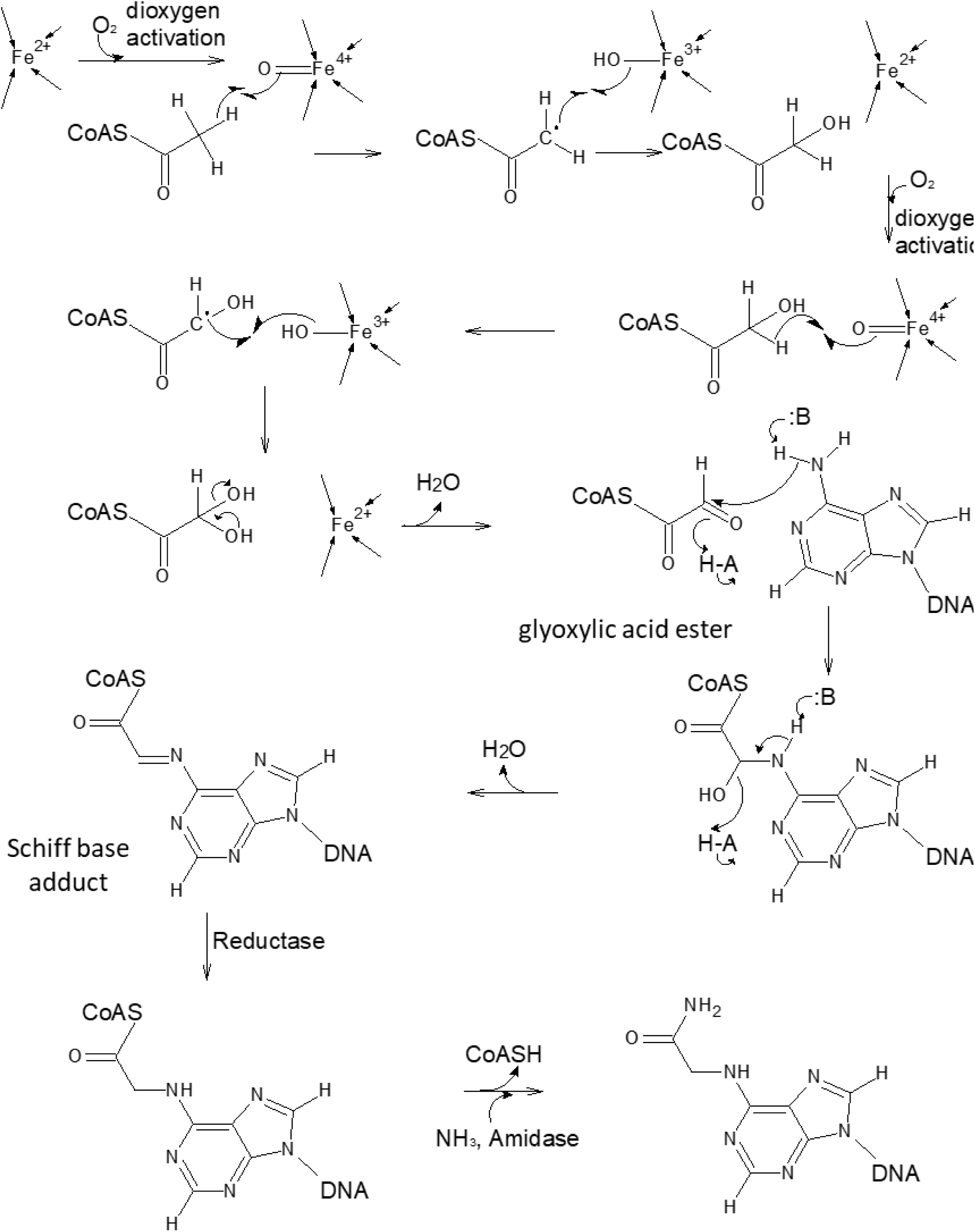
Plausible pathway for Fe^2+/3+^ ion catalyzed mediated methylcarbamoylation of adenines. Reiterative oxidation of the methyl group of acetyl CoA with molecular oxygen, catalyzed by Fe^2+^ bound in the Mom active site, generates an aldehyde group (glyoxylic acid ester) capable of forming a Schiff base with the substrate amine of adenine in DNA. Reduction of the Schiff base transfers the carboxymethyl (-CH_2_COSCoA) group to adenine. Amidation of this activated carboxymethyl group results in the final methylcarbamoyl (-CH_2_CONH_2_) modification. B-unidentified general base, HA-unidentified general acid.

### Indispensability of acetyl CoA and Fe^2+/3+^ for Mom function

Mom homologs (numerous uncharacterized proteins annotated as Mom owing to sequence similarity with phage Mu Mom) from various representative genera were aligned to identify conserved residues (Supplementary Figure S6). Alanine substitutions at conserved polar residues impaired the activity of Mom to different extents, as demonstrated by the endonuclease protection assay employed in the previous sections (Figure 5A and Supplementary Figure S7). Subsequently, each of the six functionally compromised point mutants was purified and individually tested for defects in Fe^2+/3+^ and/or acetyl CoA-binding using microscale thermophoresis assay (Figure 5B). The mutants fell into four categories based on defects in i) acetyl CoA-binding only (H48A and S114A), ii) both iron- and acetyl CoA-binding (Y49A and Y149A), iii) iron-binding only (D139A) and iv) neither iron-binding nor acetyl CoA-binding (R101A) (Figure 5B). These functionally important residues mapped to the acetyl CoA cleft in Mom, suggesting the region to be the active site of Mom (Figure 5C). Conversely, mutating a poorly conserved residue located far from the acetyl CoA cleft (e.g. L69) did not perturb Mom activity (Figures 5A and Supplementary Figure S6). Mutagenesis data further validated the relatedness between Mom and the GNAT superfamily predicted *in silico*. In GNAT members, acetyl CoA mainly contacts the protein via backbone interactions ^15^. The only few conserved side chain contacts include a tyrosine in α4 (Y149 in Mom), which stabilizes the acetyl arm of acetyl CoA, and a serine in α3 (S114 in Mom), which stabilizes the pyrophosphate arm of CoA ^14,15^. S114 and Y149 appear to furnish similar roles in Mom, evident from their conservation across Mom homologs (Supplementary Figure S6) and mutations therein impairing both acetyl CoA binding and DNA modification activity of Mom (Figures 5A to C). Surprisingly, mutation of Y149 also abolished iron binding (Figure 5B). Defects in iron-binding and DNA modification were also seen with mutations at residues Y49 and D139, which along with Y149, likely co-ordinate an Fe^2+/3+^ ion (Figures 5A-C and Supplementary Figure S7). Remarkably, the putative iron-coordinating triad maps near the acetyl group of acetyl CoA modeled within the Mom active site (Figure 5C). This explains why mutating iron-coordinating tyrosines 49 and 149 also impairs acetyl CoA binding (Figure 5B and C). The converse however is not true, with residue S114, which stabilizes acetyl CoA via the pyrophosphate arm at a region far from the active site, having no effect on iron-binding (Figure 5B and C). Interestingly, residues like Y149, long conserved for their GNAT-specific roles ^14^, seem co-opted for a novel task of iron-binding in Mom (Figure 5B).

**Figure 5.**
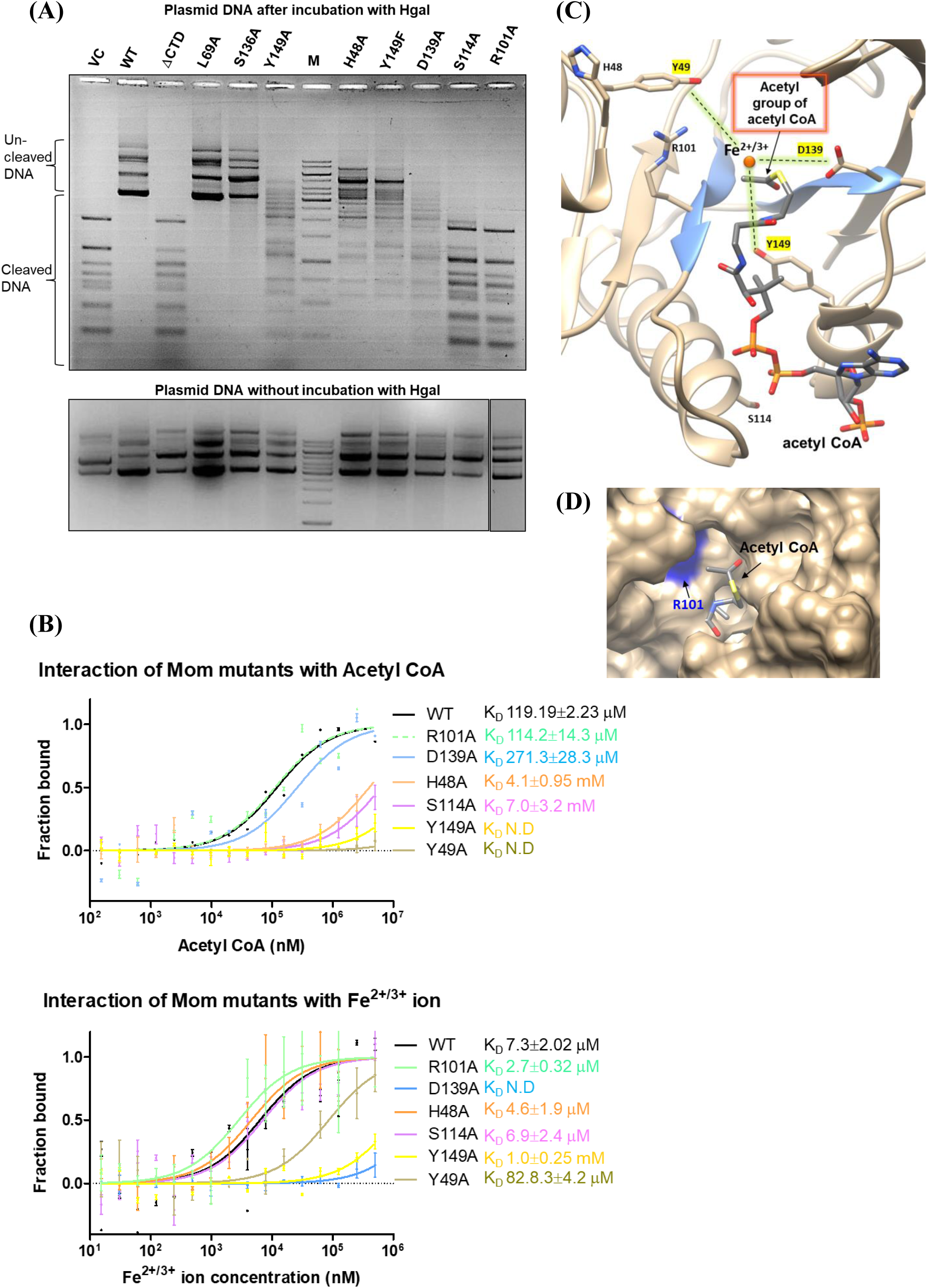
DNA modification activity and co-factor-interactions of Mom mutants. (A) Analysis of *in vivo* DNA modification activity of Mom mutants. Plasmid DNA was isolated from *E. coli* C41 cells expressing wild type or mutant Mom proteins, as indicated. VC denotes vector control and ΔCTD denotes a 175 aa long Mom construct with 56 residues from the C-terminus deleted. M represents 1kb ladder. 1 μg DNA was digested with the restriction endonuclease HgaI and products were analyzed on a 1% agarose ethidium bromide gel. Different mutations exert variable degrees of effects on the activity of Mom, with S114 and R101A compromising Mom activity most severely. Lower panel shows the undigested plasmids corresponding to those in the upper panel. **(B) Co-factor-binding defects in Mom mutants analyzed using microscale thermophoresis.** His-tagged wild-type Mom (WT) and mutants were fluorescently labeled and titrated against indicated concentrations of ligands acetyl CoA (upper panel) and Fe^2+/3+^ (lower panel). Normalized fluorescence is plotted for analysis of thermophoresis. Data were fitted using the law of mass action to determine dissociation constants (K_D_). Data shown are representative of three independent experiments. Error bars represent standard deviations of n=3 measurements. All mutants except R101A and D139A are defective in acetyl CoA binding, whereas Y49A, D139A and Y149A are compromised for Fe^2+/3+^ binding. **(C) Active site of Mom.** Residues found to be important for the DNA modification activity of Mom (side chains shown) map onto the active site pocket of Mom, with the splayed strands β4 and β5 which form the acetyl CoA-binding cleft colored blue. A Fe^2+/3+^ ion is colocalized near the acetyl arm of acetyl CoA within the active site of Mom. The tentative location of the Fe^2+/3+^ ion is depicted based on mutagenesis data, with residues critical for Fe^2+/3+^-binding highlighted. **(D) R101 residue of Mom.** Of all the point mutations introduced in the Mom active site, the effect of R101A substitution was the most detrimental to the modification activity, despite having no effect on acetyl CoA- and Fe^2+/3+^-binding. The active site cleft viewed from the acetyl end of acetyl CoA shows the acetyl group emerging into a grove, likely to be a substrate-binding cleft. The residue R101 is positioned at the opening of the cleft and appears poised for substrate recognition.

R101, a functionally critical residue in Mom is conserved amongst Mom homologs but not across the GNAT superfamily (Figure 5A and Supplementary Figure S6) ^12^. R101 maps within the active site cleft of Mom, being located near the acetyl moiety of acetyl CoA (Figure 5C and D). Surprisingly, although mutation of R101 totally abolished Mom activity (Figure 5A), binding to acetyl CoA and iron was not affected (Figure 5B). Consistent with its location in the active site and positioning near the opening of the acetyl CoA cleft and substrate-binding cleft (Figure 5D), R101 possibly is involved in substrate recognition or stabilization rather than acetyl CoA- or iron-binding. A flipped adenine base in DNA could enter the active site, placing itself near the acetyl group of acetyl CoA.

Overall, the computation-guided biochemical analyses reveal the active site of Mom as a milieu in which the acetyl group of acetyl CoA, Fe^2+/3+^ and substrate adenine come together to perform an unusual DNA modification reaction (Figure 4 and 5C). In contrast to acetyl CoA, which is well-known to occupy a conserved cleft in the GNAT fold ^14^, binding of both - iron and DNA to acetyltransferases, to our knowledge, is totally unprecedented. Our data reveals that the GNAT-like active site of Mom has retained ancestral GNAT features while gaining at least two novel functions-iron-binding and utilization of DNA as a substrate. Moreover, colocalization of iron and acetyl group in the active site of Mom supports the catalytic role of iron suggested by the depicted molecular mechanism (Figure 4).

## Discussion

While conventional chemistry fails to explain acetyl CoA as a co-factor of Mom, a radical-based mechanism can explain it, for instance, through oxidation of acetyl CoA. Also, our investigation renders several other metabolites and pathways unlikely, leaving an iron-mediated radical mechanism the most plausible one for methylcarbamoylation of adenine. Besides acetyl CoA and Fe^2+/3+^ acting together in the active site of Mom, host factors need to chip in to complete the Mom-mediated modification of DNA. Functional redundancy or essentiality in *E. coli* has precluded the identification of the host partners involved. Nevertheless, the discovery of acetyl CoA and iron in the Mom active site unravels a concrete piece in the enigmatic DNA modification puzzle. Our findings pave the way for further mechanistic studies, potential harnessing of the modification for therapeutic application of phages, and more.

Use of acetyl CoA marks a brilliant evolutionary strategy by phage Mu. Acetyl CoA is ubiquitous and abundant, with concentrations in exponentially dividing *E. coli* reaching ∼0.6 mM ^43^. Mu expresses Mom only briefly at the end of lytic infection, and cellular acetyl CoA pools are unlikely to be a limiting factor for Mom activity ^8,44^. Moreover, the central metabolic role of acetyl CoA precludes the host from dispensing with it in order to thwart DNA modification by Mom.

While simple ‘methylation’ appears to be the modification of choice for most nucleic acid modification systems, nature has, albeit less frequently, explored other more complex modifications ^33,45^. Their biological roles often extend beyond providing immunity against restriction enzymes ^1,4,6,46^ to those concerning regulation of gene expression (base J in trypanosomes ^47,48^, hydroxymethylcytosine ^49^), protein synthesis (e.g. tRNA anti-codon loop modifications ^50-53^), and structural and thermal stability of nucleic acids ^54-56^. As in the case of Mom, biosynthetic pathways for several modifications have long remained enigmatic until recently, chemistries underlying some began to be unraveled ^26,33,45,57,58^. Examples include radical SAM enzyme-mediated reactions on RNA ^26,41^ and Tet1-catalyzed hydroxylation and subsequent oxidation of methylcytosines in DNA ^40,59,60^. Notably, methylation of DNA was long thought to be too stable to be enzymatically removed, but it was subsequently found to be erasable via the action of widely occurring Tet family proteins ^40^. A property that unites extraordinary enzymes like radical SAM proteins and Tet1 dioxygenases is their ability to bind iron and catalyze free radical chemistries. Iron likely enables Mom to catalyze an unusual reaction using acetyl CoA instead of a reaction typical of the GNAT fold. We suggest that the occurrence of iron-mediated chemistry might be more common than previously thought and may be the missing piece in several other biological puzzles.

Summing up, phage Mu Mom represents evolutionary and chemical innovation at multiple levels. Mom is the first GNAT member to bind Fe^2+/3+^ and carry out a non-acylation reaction on DNA, a substrate unexplored so far by the GNAT superfamily. The novelty of DNA as a GNAT substrate is noteworthy, given that the GNAT superfamily is known to modify a vast variety of biomolecules-proteins, peptides, sugars, RNA, antibiotics, etc. ^14,15^. From an evolutionary perspective, the metal-mediated ability to generate highly reactive radical species vastly expands the catalytic capabilities of a given structural scaffold ^39^. Thus, characterization of Mom unveils not only the catalytic versatility possible within the GNAT world, but also a novel class of enzymes whose chemistry lies outside the canonical borders of the GNAT superfamily. Evolutionary tinkering of enzyme active sites thus explains how a disproportionately high number of functions evolve from a small number of folds ^61^. As for phage Mu, the lucky evolutionary accident of rebooting an ancestral enzyme fold enables it to successfully penetrate the anti-phage arsenal of its bacterial hosts.

## Materials and Methods

The materials and methods used in this study are described in detail in Supplementary data. Information includes expression and purification of Mom, *in vivo* activity assays, structure and co-factor prediction, ligand interaction analyses using microscale thermophoresis and differential scanning fluorimetry, mass spectrometric analysis of the Mom-modified nucleoside, ferrozine assay to detect iron binding, transposon mutagenesis screens and affinity based pull-down assays for Mom-host factor interaction, etc.

## Acknowledgements

This research was funded by various grants to V.N. from the Department of Science and Technology, Indian Institute of Science-Department of Biotechnology partnership, Jawaharlal Nehru Centre for Advanced Scientific Research-Department of Biotechnology partnership and J.C. Bose Fellowship to V.N. S.K. was supported by fellowship from the Council of Scientific and Industrial Research, India. We are grateful to Karl Drlica for critical reading of the manuscript, and thank P. Balaram, K. R. Prasad, Debasisa Mohanty, Amrita Hazra, J. Gowrishankar, D.N. Rao, Ujjwal Rathore, Souvik Bhattacharyya and members of the Nagaraja laboratory for scientific input and discussions. We thank Ashish Rangra, Samata, and Rahul for technical help, Shubhada for bioinformatics support, Sunita and Srilatha from the Department of Biotechnology-supported mass spectrometry and biacore facilities, respectively, at the Indian Institute of Science for technical assistance, and Saji Menon and Sivaramaiah Nallapeta of Nanotemper technologies for their assistance with microscale thermophoresis and nanoDSF measurements.

