## Supplementary Materials methods, figures and tables for "Emergence of a novel immune-evasion strategy from an ancestral protein fold in bacteriophage Mu"

**Supplementary Materials and Methods**

***Bacterial strains and plasmids***

The *E. coli* strains and primers used in the study are described in Supplementary data Table 4 and 5, respectively.

***Purification of Mom***

All chemicals used in the study were procured from Sigma, unless otherwise stated. *E. coli* C41 ($F^-$ *ompT hsdS*$_B$ ($r_B^-$ $m_B^-$) *gal dcm* (DE3)) (laboratory stock) was used for the overexpression of Mom. The *mom* operon was cloned in the BamHI site of pVN6, a modification of the T7 expression vector pET11d (1, 2). A regulatory plasmid pNC1, derived from pLysE, served to express T7 lysozyme and to minimize leaky expression (1). About 10 g cell pellet (*E. coli* C41 harbouring the plasmids pNC1 and pVN6*mom*) was resuspended in 30 ml lysis buffer (10 mM HEPES-NaOH (pH 7.4), 14.33 mM β-mercaptoethanol, 1 M NaCl, 1 mg/ml lysozyme and 1 mM PMSF) and sonicated on ice using a Sonics macrotip set at 36 % amplitude, 20 pulses of 2 s each). The sonicated extract was centrifuged for 2 h at 100,000g. Nucleic acids were precipitated using 1% (v/v) polyethyleneimine. Proteins in the supernatant fluid were fractionated using ammonium sulfate (3). A series of cation exchange resins, including phosphocellulose (Whatman), heparin sepharose and sulfopropyl sepharose (GE), were employed to achieve a near homogenous preparation of Mom. Mom-containing fractions were dialyzed against storage buffer (10 mM HEPES-NaOH (pH 7.4), 14.33 mM β–mercaptoethanol, 300 mM NaCl) and concentrated using Amicon ultra centrifugal filter with 10 kDa cut-off (Millipore). Aliquots were flash-frozen in liquid nitrogen and stored at -80°C. Purity and concentration of Mom were determined by densitometry analysis from SDS-PAGE using standard proteins of known concentrations.

For purification of MBP (Maltose binding protein) and MBP-tagged Mom, cell extracts were prepared from *E. coli* C41 harbouring the pMALC2x plasmid (for expressing MBP) and pBAD*mom*-MBP (MBP-Mom fusion cloned in pBAD24 and induced using arabinose) and bound to amylose resin (NEB). The beads were washed with 20-30 volumes of column buffer (10 mM HEPES pH 7.4, 10 mm β-mercaptoethanol, 200 mM NaCl) and eluted with one column volume of column buffer containing 10 mM maltose. Proteins were finally dialyzed against column buffer and stored at -80°C.

***Glutaraldehyde cross-linking of protein***

Purified Mom was dialyzed against buffer containing 50 mM sodium phosphate (pH 7.6), 14.33 mM β-mercaptoethanol, 0.1 mM EDTA and 150 mM NaCl. 25% (v/v) glutaraldehyde was thawed on ice and diluted in MilliQ water (EMD Millipore) immediately before use. In a 20 μl reaction volume, 2 μg Mom was incubated with glutaraldehyde (0.0025%-0.025%) for 5 min at room temperature. The reactions were stopped using 0.4 M Tris-HCl (pH 8.0) and analyzed by 13% SDS PAGE.

***Far-UV Circular Dichroism (CD)***

CD spectra were recorded on a Jasco J-715C spectropolarimeter flushed with nitrogen gas. 3.33 μM Mom in 1X PBS (phosphate buffered saline) was subjected to measurements using a 1 mm path length quartz cuvette, with a scan rate of 50 nm/min, a response time of 4 s and a bandwidth of 2 nm. Each spectrum was an average of three scans. Buffer spectra were also acquired under similar conditions and subtracted from protein spectra before analysis.

***Fluorescence Spectroscopy***

All fluorescence spectra were recorded at 25°C on a SPEX Fluoromax3 spectrofluorimeter. 2 μM Mom in 1X PBS was subjected to excitation at 280 nm and emission was recorded from 300 to 400 nm. For measuring the fluorescence of the denatured Mom, the protein was denatured using 6 M guanidine HCl (pH 7.4). Each spectrum was an average of three consecutive scans. Buffer spectra were also acquired under similar conditions and subtracted from protein spectra before analysis. All fluorescence experiments were carried out in buffer $A_{600}$ (20 mM Tris-HCl (pH 7.4) at $4^0$C, 0.1 mM EDTA, 14.33 mM β–mercaptoethanol, 10% glycerol, 350 mM NaCl).

***Surface plasmon resonance (SPR)***

All SPR experiments were performed with a Biacore 2000 (Biacore, Uppsala, Sweden) optical biosensor at 25°C. In the assay, biotinylated 32 bp double stranded DNA was attached to the surface of a research-grade streptavidin chip (GE Healthcare). The binding of Mom to this surface was examined. A sensor surface without DNA served as a negative control for each binding interaction. Various concentrations of Mom, serially diluted in 10 mM HEPES-NaOH (pH 7.4), 300 mM NaCl and 14 mM β-mercaptoethanol was run across each sensor surface at three different concentrations (2.24 nM, 4.48 nM and 8.96 nM) in a running buffer of the same composition as the dilution buffer with the addition of 0.005% Tween 20.

***Analysis of in vivo activity of Mom***

*E. coli* C41 cultures harboring the regulatory plasmid pNC1 and the wild type or mutant *mom* plasmids were grown at 37°C to mid exponential phase and induced with 0.5 mM IPTG. Incubation was continued for 12 h at 16°C. Alternately, when using the pBAD expression system for *mom* expression, *E. coli* cultures harbouring the wild type or mutant *mom* plasmids were grown to mid log phase and induced with 0.2% arabinose. Incubation was continued overnight at 37°C. Plasmid DNA was isolated from the cultures using Qiagen miniprep kits and 1 μg DNA was digested with HgaI (2 units/μl, NEB). Products were analyzed on a 1% agarose ethidium bromide gel.

***Investigating the interacting partner(s) of Mom***

MBP-Mom fusion protein or MBP control were expressed in *E. coli* C41 from arabinose inducible pBAD promoter. Cells were grown in 600 ml LB medium in the presence of 0.2% glucose till $O.D_{600}$ reached 0.4, after which the culture was induced with 1% arabinose for 1h. Cells were resuspended in column buffer (20 mM Tris-HCl (pH 7.4), 10 mM β-ME, 25 mM NaCl) and sonicated. Clarified cell extract was applied to 1 ml amylose resin (NEB) pre-equilibrated with column buffer. The resin was washed with 5 ml column buffer containing 200 mM NaCl followed by elution with column buffer containing 10 mM maltose. Samples were briefly electrophoresed on 12% SDS PAGE, stained with Coomassie blue and submitted for protein identification at Alphalyse, Denmark. The service included in-gel trypsin digestion of proteins followed by injection of concentrated peptides on a Dionex nano-LC system and MS-MS analysis on a Bruker Maxis Impact Q-TOF instrument.

***Generation of Mom mutants***

Plasmid pVN6*mom* was used as a template for the generation of *mom* mutants. Mutations were introduced by site-directed mutagenesis using inverse PCR with adjacent non-overlapping primers (4). Forward primers harbored the desired mutation. Primer sequences are listed in Supplementary data Table 5.

***Mass spectrometric analysis of Mom***

100 ng pure Mom protein in 5 mM HEPES-NaOH (pH 7.4), 150 mM NaCl and 7.16 mM β-mercaptoethanol was injected on a C18 reversed phase column connected to an ESI Maxis Impact mass spectrometer and eluted using a gradient of acetonitrile. The mass detected by mass spectrometry was compared to the theoretical mass calculated for Mom.

***HPLC and mass spectrometry of Mom modified deoxyadenosine***

Genomic DNA from *E. coli* C41 cells overexpressing Mom was isolated using a Qiagen kit. About 200 μg of genomic DNA was digested with 5 units of DNase I (NEB) at $50^0$C for 5 h. The di-, tri and oligonucleotide products generated by DNase I were digested to mononucleotides with 0.01 units (1 mg) of exonuclease Phosphodiesterase I from *Crotalus adamanteus* venom (Sigma) overnight at $37^0$C. Mononucleotides were dephosphorylated with 1 unit of bacterial alkaline phosphatase (Sigma) for 4 h at $37^0$C. High molecular weight contaminants such as proteins or undigested nucleic acids, if any, were removed by passing the digestion reaction through a 3 kDa centricon membrane (Millipore). Standards were prepared by dephosphorylating dATP, dGTP, dTTP and dCTP with bacterial alkaline phosphatase. Nucleosides were resolved on an Acclaim 120 C18 column (4.6 x 150 mm column, 3 μm, Dionex) eluted at 0.3 ml/min with 5 mM ammonium acetate pH 6.0 using an acetonitrile (0 to 40%) gradient on a Dionex Ultimate 3000 HPLC system (Thermo) (Supplementary data Table 6). Individual peaks were collected and injected into a Bruker Daltonics ESI-MS ion trap mass spectrometer.

***Analysis of thermostability of Mom in the presence of ligands***

Purified Mom (12 μM) in HEPES buffer (10 mM HEPES-NaOH (pH 7.4), 14.33 mM β-mercaptoethanol, 100mM or 300 mM NaCl) was incubated with ligand for 5 min on ice followed by 10 min at room temperature. Reactions (10 μl each) were set up in duplicates, loaded into nanoDSF grade glass capillaries and placed into Prometheus NT.48 (NanoTemper Technologies). A linear thermal ramp (1°C/min, 20-80°C) was applied and the thermal unfolding of Mom was monitored by detecting the temperature-dependent change in tryptophan fluorescence at emission wavelengths of 330 and 350 nm. Protein melting points ($T_m$) were calculated from the first derivative of the ratio of tryptophan emission intensities at 330 and 350 nm by the Prometheus NT.Control software.

***Binding affinity quantifications using* microscale thermophoresis *(MST)*** Affinity measurements of purified Mom with various ligands were carried out on a Monolith NT.115 instrument (NanoTemper Technologies GmbH). Mom was fluorescently labeled using NanoTemper's Protein Labeling Kit RED-NHS or RED-tris-NTA dye (for his-tagged Mom and mutants). Ligands including acetyl CoA, S-adenosyl methionine (SAM), coenzyme A (CoASH), malonyl coenzyme A and salts of various metal ions were resuspended or dissolved in MST buffer (10 mM HEPES-NaOH (pH 7.4), 300 mM NaCl and 14 mM β-mercaptoethanol) to prepare a 16 step two-fold dilution series. Diluted ligands were incubated with 50 nM of labeled Mom at room temperature for 5 min and reactions were spun at 17,000g for 5 min. Supernatants were loaded into NanoTemper premium coated capillaries. MST measurements were performed at $25^0$C using 60% MST power and 90% LED power. Data analyses were carried out using NanoTemper analysis software.

***Analytical Gel Filtration***

Mom was subjected to size exclusion chromatography using an analytical Superdex 75 column (GE Healthcare, column volume $V_t$= 24 ml, void volume = 8 ml) on an AKTA purifier FPLC system (GE). Briefly, pure fractions of Mom protein eluted from heparin sepharose column were injected into the gel filtration column equilibrated in 50 mM sodium phosphate (pH 7.4), 0.1 mM EDTA, 14.33 mM β–mercaptoethanol and 800 mM NaCl. 50 μg of Mom was loaded on to the column and eluted at a flow rate of 0.25 ml/min. Calibration curve was obtained using molecular weight calibration markers (GE).

***Ferrozine assay***

Mom, MBP-Mom fusion and MBP proteins were dialyzed against 10 μM ferrous ammonium sulfate (FAS) in HMN buffer (10 mM HEPES-NaOH (pH 7.4), 1 mM β-ME, 200 mM NaCl) for 1.5 h. Free iron was removed by dialysis against HMN buffer. Proteins were subjected to Ferrozine assay to quantitate bound iron (5). Intrinsically bound $Fe^{2+/3+}$ was quantified by directly subjecting 70 μg MBP-Mom fusion protein to Ferrozine assay (5).

***In silico prediction of structure and co-factor of Mom***

The amino acid sequence of Mom was submitted to various structure prediction servers such as Robetta (6) and I-TASSER (7) available online at http://www.robetta.org/submit.jsp, http://zhanglab.ccmb.med.umich.edu/I-TASSER/. The modeled structures of Mom were submitted to the COFACTOR server available online at http://zhanglab.ccmb.med.umich.edu/COFACTOR/ in order to model the Mom-acetyl CoA complex, obtain functional insights including ligand-binding site, gene-ontology terms and enzyme classification derived from the best functional homology template. Models were visualized and analyzed using The PyMOL Molecular Graphics System, Version 2.0 Schrödinger, LLC. or the UCSF Chimera software available online at http://www.cgl.ucsf.edu/chimera (8). The structure of the N- and C-termini of Mom is difficult to reliably predict owing to lack of counterparts in template structures and so residues 1-28 and 195-231 have not been depicted in the model (9).

***Multiple sequence alignment of Mom homologs***

Representative homologs of Mom obtained using an NCBI-BLASTp search (Altschul 1990) were aligned using Constraint-based Multiple Alignment Tool from NCBI and the alignment was visualized using Boxshade.

***Transposon insertion library preparation and Screening for modification deficient mutants***

*E. coli* TG1 or MG1655 were transformed with pBAD *mom* operon (with his-tagged *mom*) and a single colony was inoculated in 2 ml TBMM (1% tryptone, 0.5% NaCl, 10 mM $MgSO_4$, 0.2% maltose) and grown overnight or for 6 h or till $O.D_{600}$ ~ 0.4 at $37^0$C. For the *thi*⁻ strain BW25113, 2μg/ml thiamine was supplemented. The culture was concentrated 10 times by centrifugation at 4000g for 10 min at $4^0$C and resuspending in LB (1% tryptone, 0.5% NaCl, 0.5% yeast extract). To 100 μl suspension, 20 μl phage lambda NK1316 was added. The multiplicity of infection was between 0.1 and 1. Incubation was carried out for 15 min at room temperature followed by 15 min at $37^0$C. The mixture was transferred into a 50 ml polypropylene tube and 5 ml LB containing 50 mM sodium citrate was added. The free phage was washed by spinning the cells at 4000g for 10 min at $4^0$C and discarding the supernatant. An additional wash with LB-citrate was given. Steps 4-5 were repeated once more. Note: Removal of free phage is not necessary. The culture was grown for 30 min-1 h at $37^0$C with shaking at 180 rpm. An aliquot of 50 μl was plated directly on LB containing 100 μg/ml ampicillin, 30 μg/ml kanamycin and 0.2% glucose to estimate the library size. The remaining culture was spun down and resuspended in 100 μl fresh LB and plated on selective media containing 1% arabinose. Survivors, or colonies that grew on 1% arabinose media, were patched on 0.2 and 1% glucose agar and simultaneously inoculated in LB or terrific broth containing 0.2% glucose and appropriate antibiotics. Colonies that continued to grow upon sub-culturing were inoculated in 5 ml LB and grown till log phase and induced with 1% arabinose. Plasmids were extracted using Qiagen miniprep kit and tested for the presence or absence of Mom-mediated DNA modification. Plasmids isolated from the survivors were introduced into BW25113 strain to assay for intactness of *mom* function.

***Protocol for λNK1316 phage lysate preparation***

*E. coli* LE392 culture was grown to saturation in TBMM. Serial dilutions ($10^{-1}$ to $10^{-7}$) of λNK1316 phage lysate were made in TMG buffer and 20 μl mixed with 200 μl of LE392 cells (10). The mix was incubated at room temperature for 10 min followed by 20 min at $37^0$ C and added to 3 ml top agar. The agar was mixed well and poured on top of basal TBI agar. Plates were incubated overnight at $37^0$C. A single plaque was picked with a micropipette and transferred into a 50 ml flask containing 10 ml LB supplemented with 10 mM $MgSO_4$ and 0.1 ml freshly grown overnight culture of LE392. The flask was shaken at $37^0$C for 4-5 h, after which the cultures became cloudy and then clear. A few drops of chloroform were added and the mixture vortexed well for 30 s. The mixture was allowed to sit for 10 min followed by spinning at 2500g for 10 min at $4^0$C. The supernatant was recovered and stored at $4^0$C.

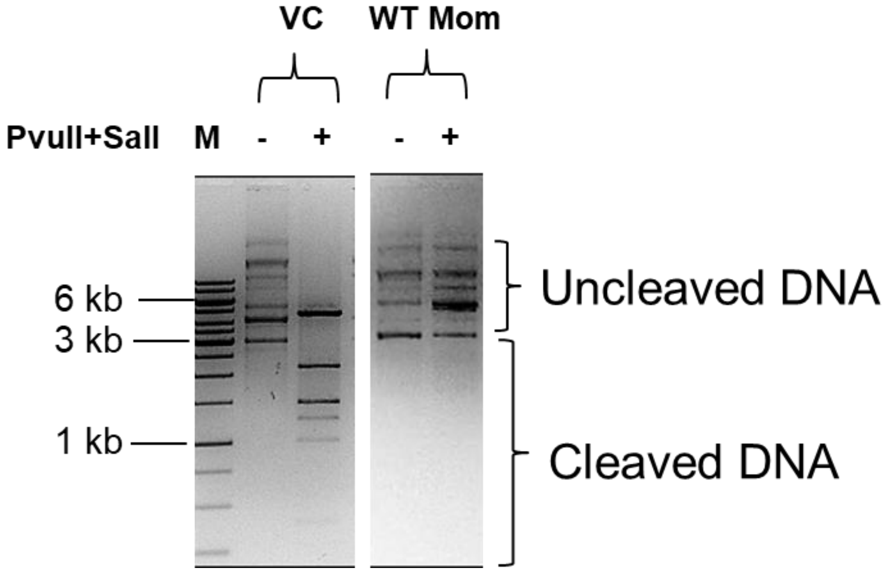


**Supplementary Figure S1. Analysis of *in vivo* activity of Mom using PvuII and SalI restriction endonucleases.** Plasmid DNA was isolated from *E. coli* C41 cells expressing wild type Mom or empty vector (vector control, VC). 1 μg DNA was digested with the restriction endonucleases PvuII and SalI and products were analyzed on a 1% agarose ethidium bromide gel. While DNA from Mom expressing cells is resistant to PvuII and SalI, DNA from vector control cells is sensitive to the enzymes.

Karambelkar et al., Supplementary Figure S2

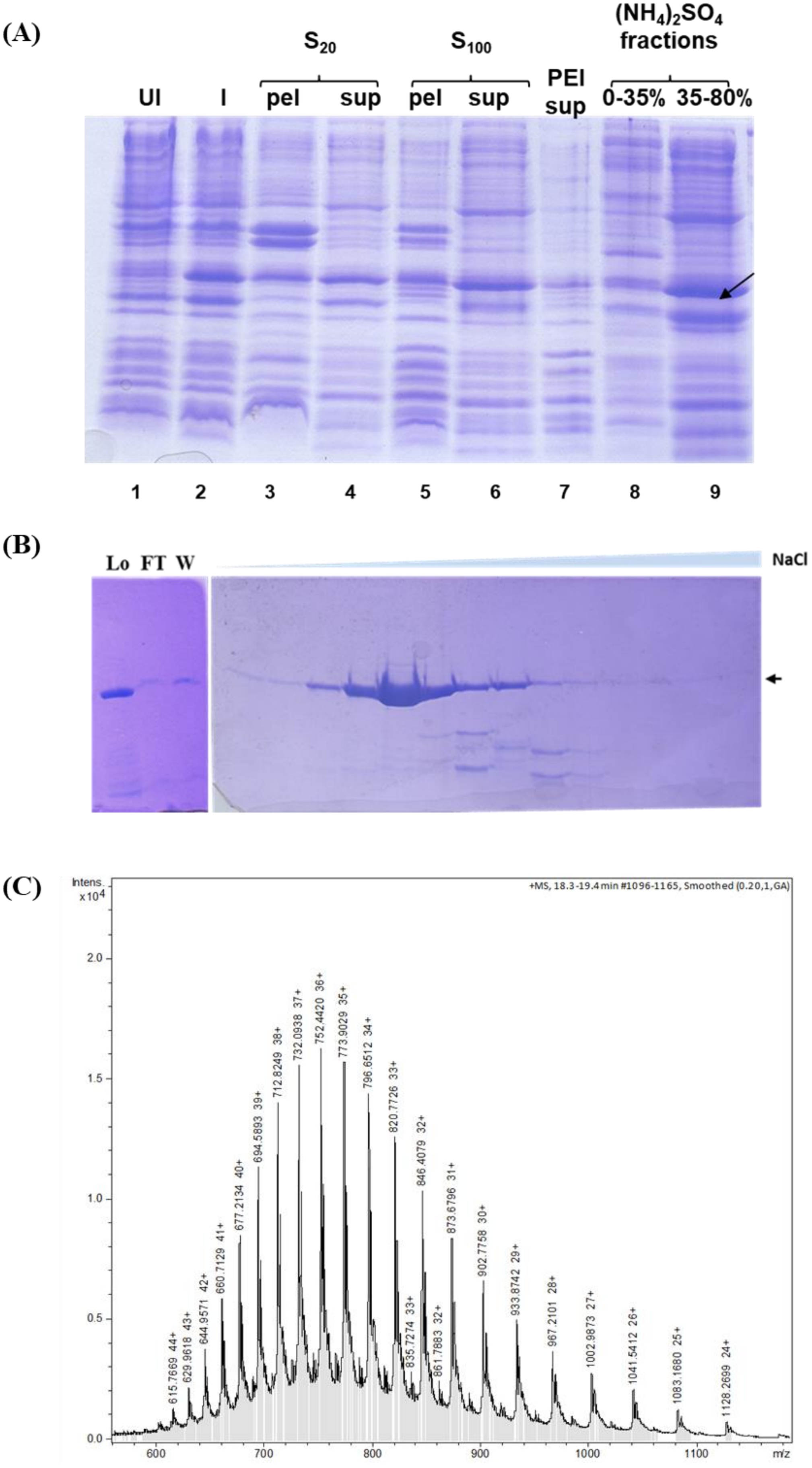

Karambelkar et al., Supplementary Figure S2

(D)

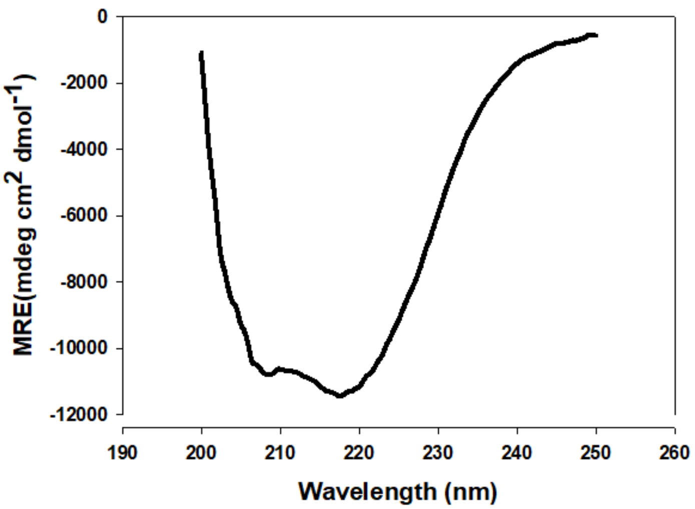


(E)

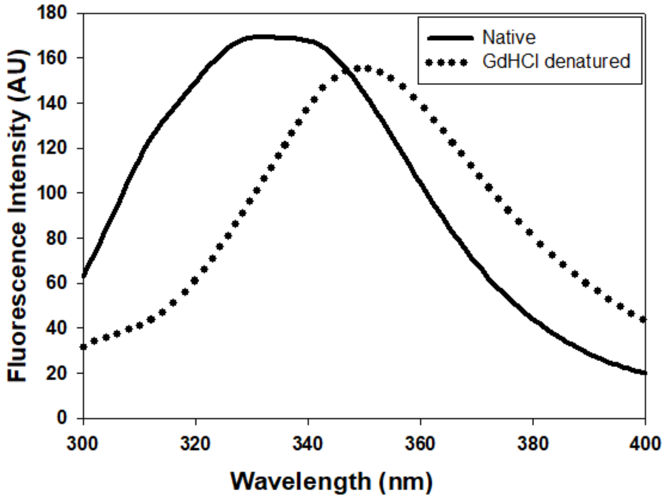


(F)

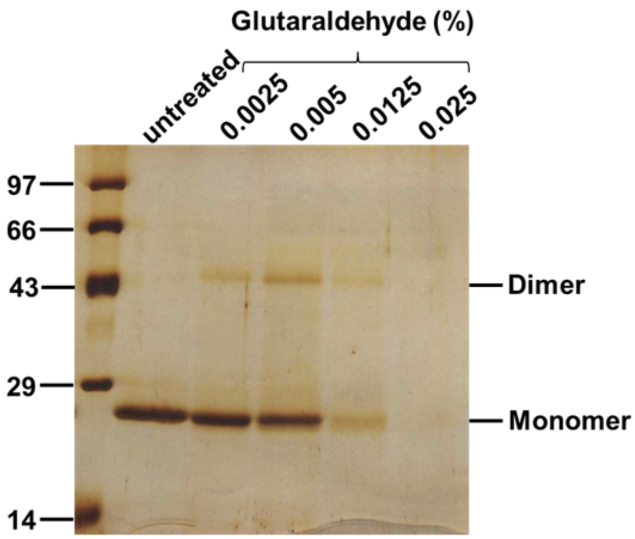

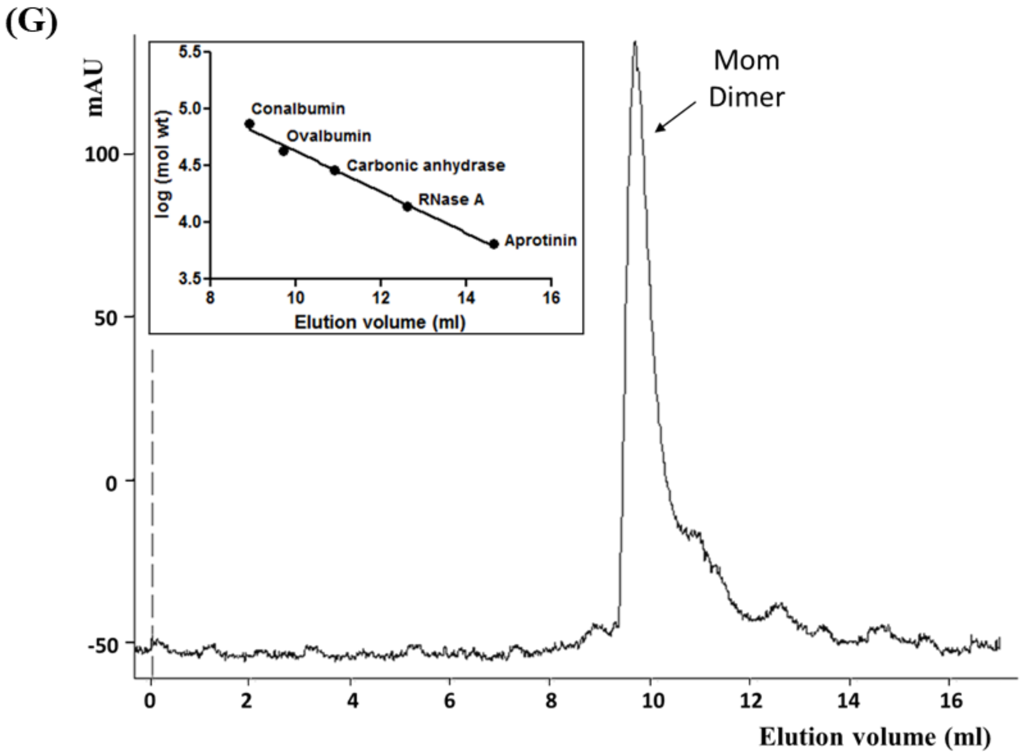


**Supplementary Figure S2. Isolation and characterization of Mom. (A) Expression and purification of Mom.** Mom expressing cultures were sonicated and $S_{100}$ cell extracts were fractionated using PEI and ammonium sulfate. UI; uninduced, I; induced, pel; pellet, sup; supernatant, $S_{20}$ and $S_{100}$; supernatants of 20,000g and 100000g centrifugation steps, respectively. The position of Mom has been indicated by an arrowhead. **(B) Sulfopropyl sepharose (SP) chromatography for Mom purification.** Protein-enriched fractions from heparin sepharose were loaded on SP sepharose column. Elution was carried out by employing an NaCl gradient (300 mM to 1M). Fractions were analyzed on a 15% SDS PAGE gel. Lo; load, FT; flow-through, W; wash. Arrow indicates the position of Mom. **(C) Mass spectrometric analysis of Mom.** Purified Mom protein was injected on a C18 reversed phase column connected to an ESI Maxis Impact mass spectrometer and eluted using a gradient of acetonitrile. The mass of Mom (27050.9372 Da) detected by ESI-MS is identical to the expected mass. **(D) Far-UV CD spectrum of Mom.** The spectrum was buffer-corrected and was obtained at 25 °C with 3.33 µM Mom in buffer containing 20 mM Tris-HCl pH 7.4, 100 mM NaCl and 2 mM β-mercaptoethanol, with a 0.1-cm path length cuvette, a scan-rate of 50 nm/min, a response time of 4 s and a bandwidth of 2 nm. Data reported are averaged over three scans. MRE, mean residue ellipticity. **(E) Fluorescence emission spectrum of Mom.** The spectrum was obtained at 25 °C with a final protein concentration of 2 μM in PBS, pH 7.4 (solid line), or in the presence of 6 M guanidine hydrochloride (dashed line). The excitation wavelength was 280 nm, and emission was recorded from 300 to 400 nm. A.U. denotes arbitrary units. **(F) Glutaraldehyde cross-linking of Mom.** Purified Mom (2 μg) was incubated with indicated concentrations of glutaraldehyde and the cross-linked products were analyzed on a 13% SDS-PAGE. The sizes of molecular weight markers in kDa are indicated. **(G) Analytical gel filtration analysis of Mom.** Gel filtration elution profile of Mom on an analytical Superdex 75 column is shown. The absorbance at 220 nm is plotted as a function of the elution volume. Mom (27 kDa) elutes at 9.6 ml, a position expected for a 50 kDa protein, indicating a dimeric state. The inset shows the calibration curve using the marker proteins conalbumin (75 kDa), ovalbumin (43 kDa), carbonic anhydrase (29 kDa), ribonucleaseA (13.7 kDa), aprotinin (6.5 kDa).

**Karambelkar et al., Supplementary Figure S3**

**(A)**

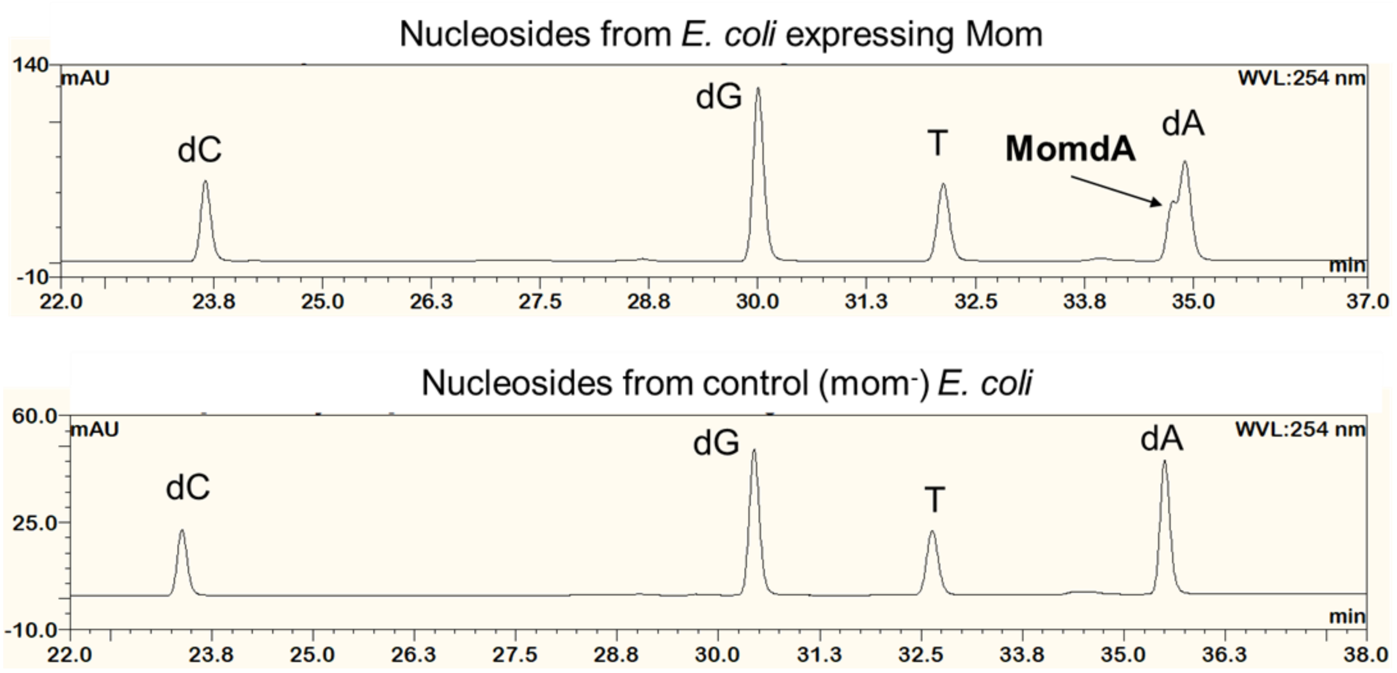

Karambelkar et al., Supplementary Figure S3

(B)

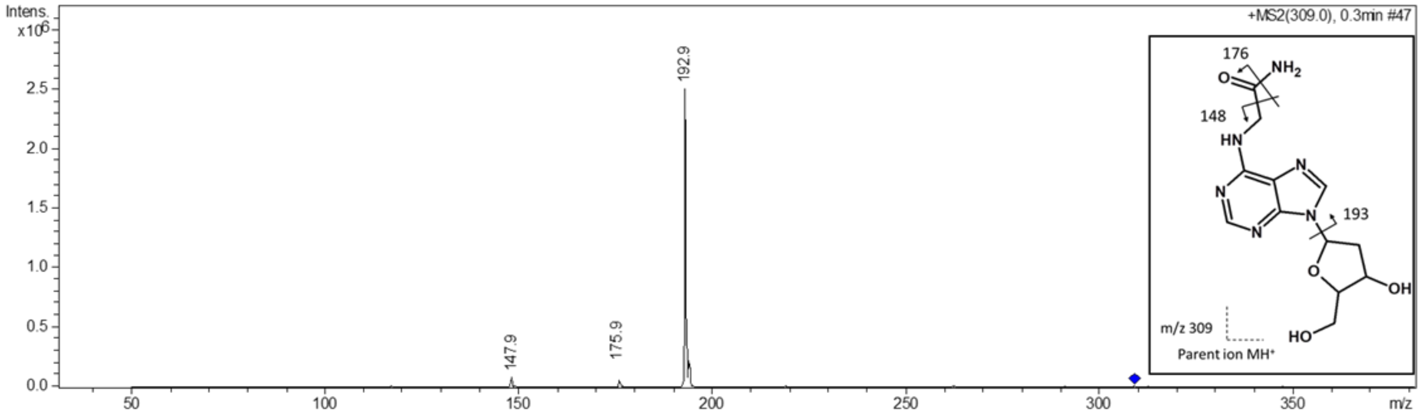


(C)

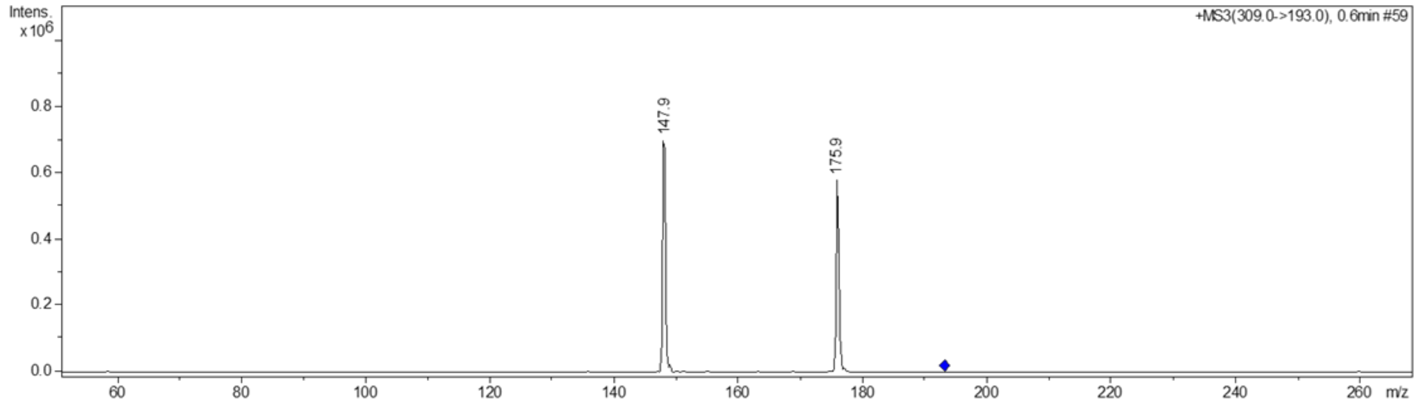


(D)

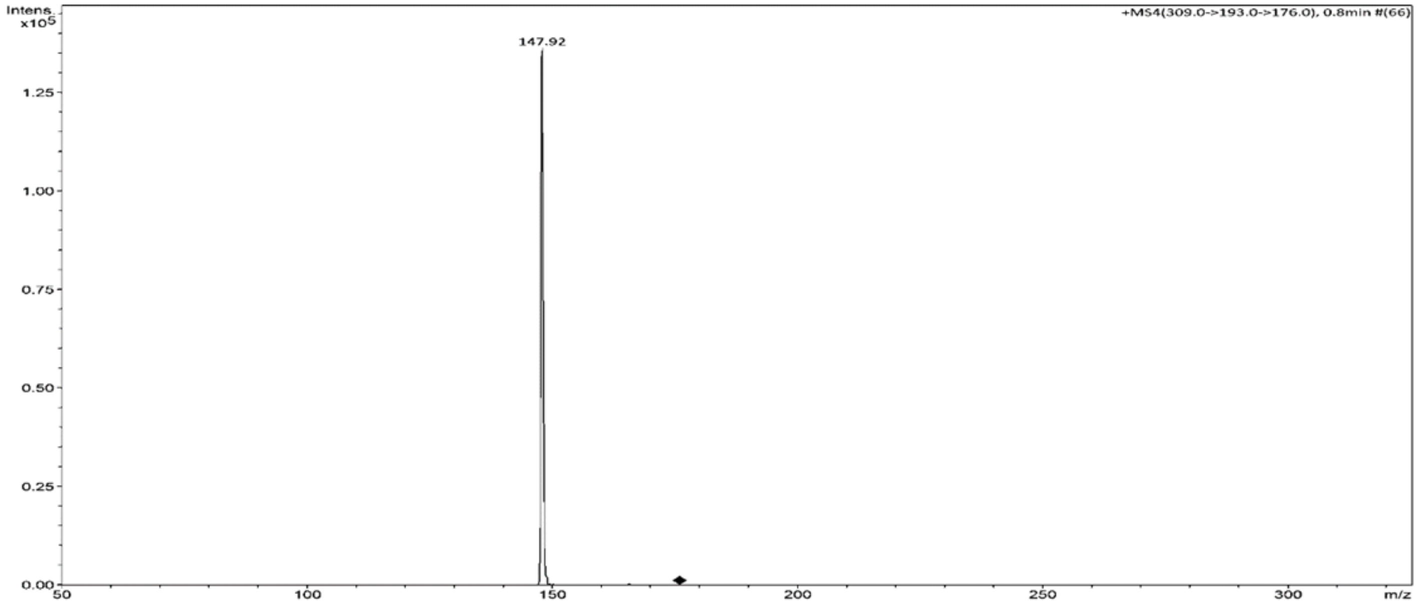

**Supplementary Figure S3. Mass spectrometric analysis of Mom-modified deoxyadenosine. (A) RP-HPLC analysis of nucleosides from Mom expressing cells.** Genomic DNA digests from *E. coli* cells expressing Mom (upper panel) and those harboring the empty vector (lower panel) were chromatographed on a C18 reversed-phase HPLC column using a gradient described in Supplementary Table 6. Mom-modified nucleoside MomdA elutes just before unmodified deoxyadenosine, as indicated by the arrowhead, and is absent in the control sample. **(B) Collision induced dissociation of momdA parent ion.** MomdA, the nucleoside uniquely present in *mom* expressing cells, was subjected to successive rounds of collision-induced dissociation to analyze its structure. An m/z of 309 for momdA parent ion corresponded to monoprotonated methylcarbamoyldeoxyadenosine. The parent ion yielded a major product of m/z 193, expected from cleavage of the glycosidic bond, and two minor products of m/z 176 and 148, expected to result from the fragmentation pattern indicated in the inset. **(C) Fragmentation profile of the product ion m/z 193, arising from fragmentation of momdA parent ion of m/z 309. (D) Fragmentation profile of the product ion m/z 176, arising from fragmentation of ion of m/z 193.**

**Karambelkar et al., Supplementary Figure S4**

(A)

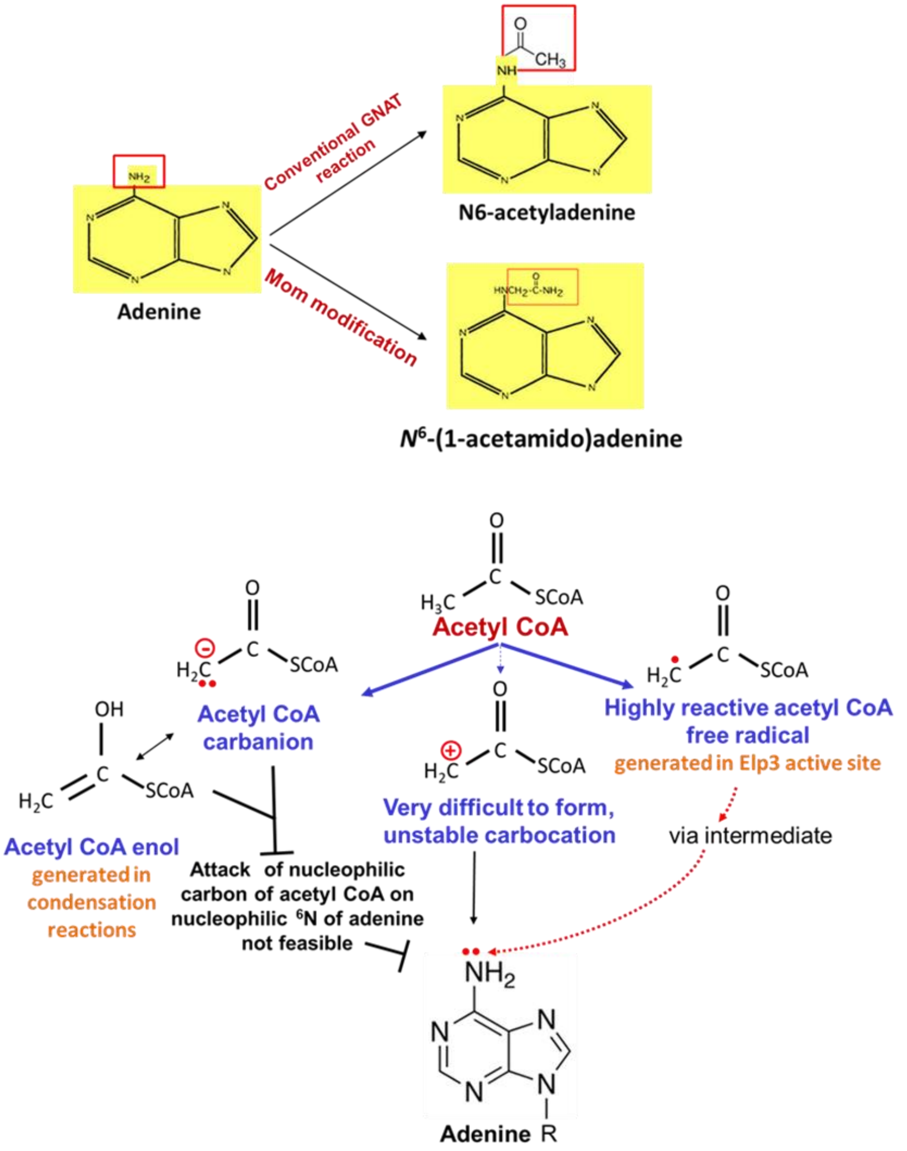


(B)

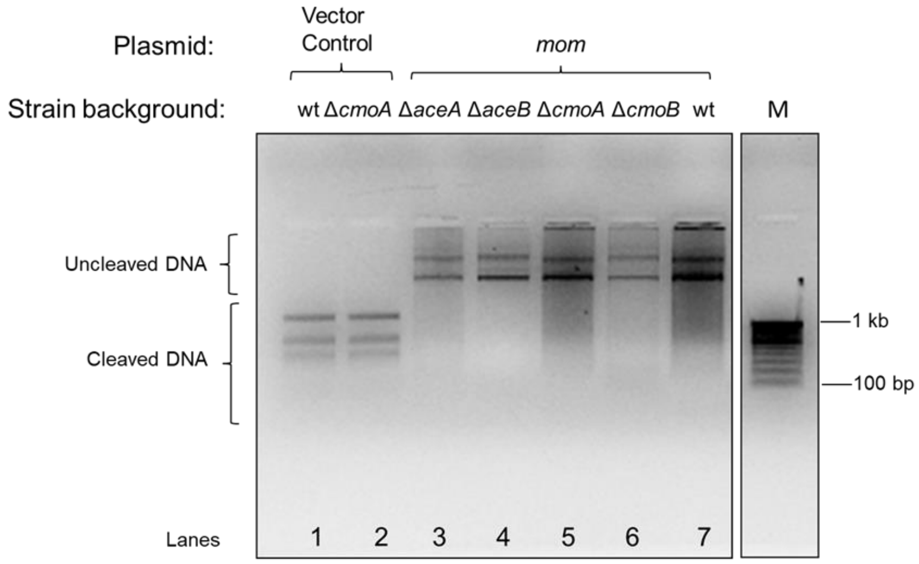

**Karambelkar et al., Supplementary Figure S4**

**(C)**

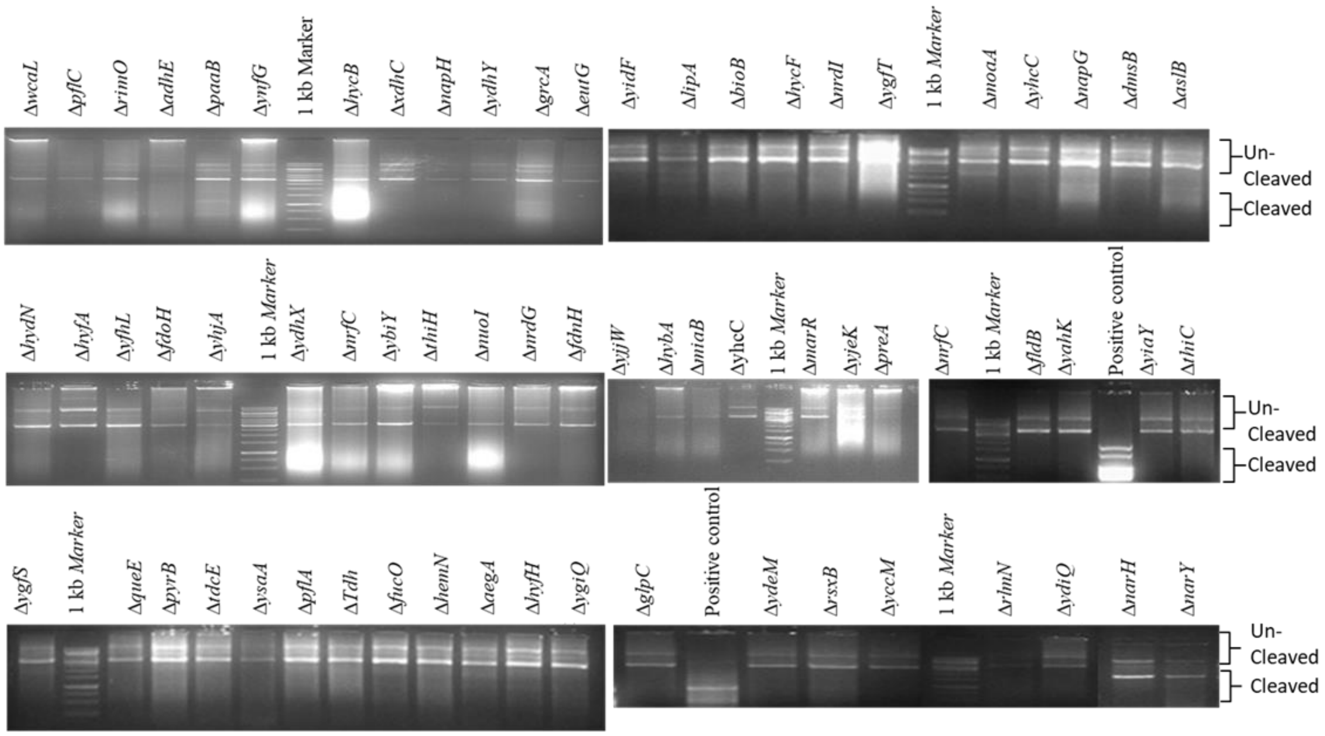


**(D)**

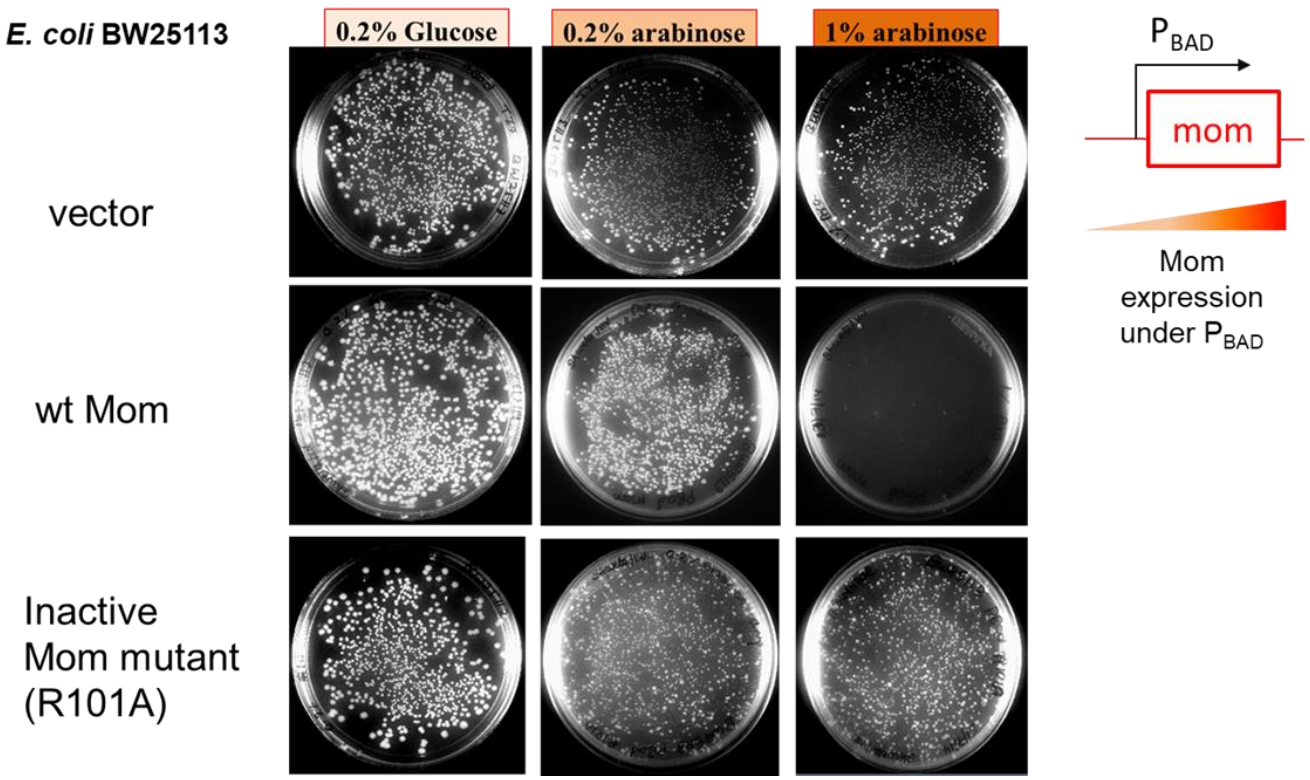

**Karambelkar et al., Supplementary Figure S4**

**(E)**

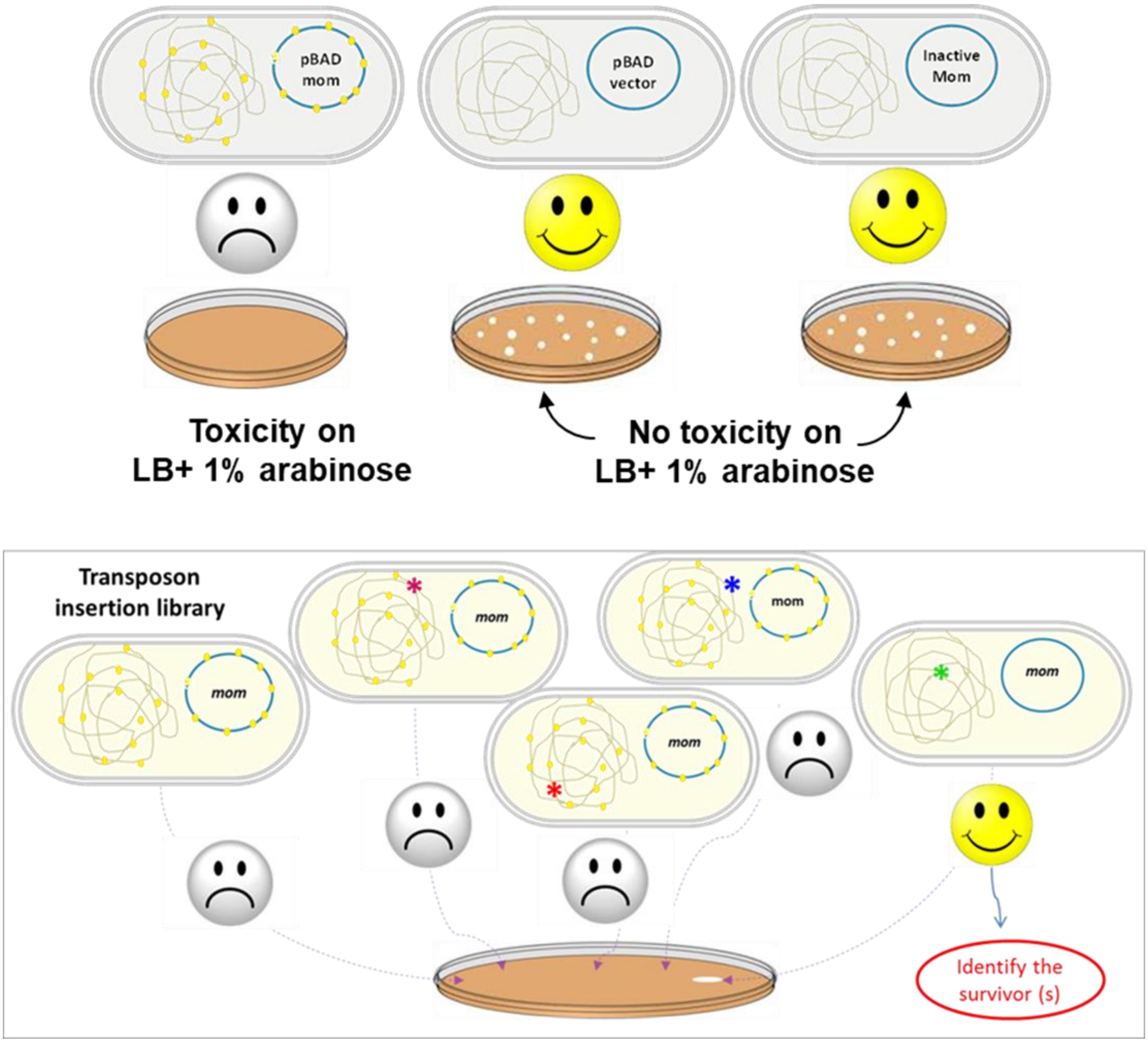

**Supplementary Figure S4. Candidate-based and genome-wide approaches to identify host genes involved in *mom* activity.**

**(A) Conventional and unconventional chemistries known for acetyl CoA.** Classic GNAT reaction involves the carbonyl carbon of acetyl CoA and leads to the transfer of an acetyl group on the free amino group of the substrate (11). However, GNAT mechanism cannot explain Mom modification, which involves the transfer of a methylcarbamoyl group instead of acetyl group. Apart from GNAT mechanisms, acetyl CoA is known to carry out other chemistries. Condensation reactions involve generation of a carbanion on the methyl group of acetyl CoA (12). On account of being nucleophilic, the carbanion cannot attack nucleophilic acceptors like the $N^6$ amino group of adenine. The acetyl CoA carbocation depicted in the center is energetically unstable and has not been observed in nature, rendering it an unlikely possibility. The newest chemistry known for acetyl CoA is one that occurs in the active site of the radical SAM protein Elp3. Elp3 forms an acetyl CoA radical as depicted, which can be conceived to attack adenine via an intermediate, if Mom employs a free radical-based mechanism (13). **(B) The glyoxylate pathway and carboxy-SAM pathway are not involved in Mom modification.** *E. coli* BW25113 and isogenic strains knocked out for glyoxylate pathway (*aceA*, *aceB*) and carboxy-SAM pathway (*cmoA*, *cmoB*) were transformed with either vector control (VC) plasmid (lanes 1 and 2) or pBAD *mom* plasmid (lanes 3 to 7). Transformants were inoculated in LB and grown till mid log phase followed by overnight induction at 37 °C with 0.2% arabinose. Plasmids were extracted and subjected to digestion with HgaI, which cannot digest Mom-modified DNA but can cleave unmodified DNA. **(C) *In vivo* activity assay of Mom in various single gene knock-out backgrounds.** For investigating the role of radical SAM (rSAM) mechanism in Mom catalyzed modification, 67 different single gene knock-out strains for *E. coli* were procured from the Keio collection (Coli Genetic Stock Center, Yale University) and are listed in Supplementary Table 3. The indicated knock-outs are deleted for genes encoding proteins with predicted or experimentally validated iron sulfur (FeS) clusters typical of rSAM proteins. If any of these rSAM/Fe-S genes are involved in the Mom modification pathway, then Mom modification cannot occur in the cells deleted for the specific gene(s). The plasmid pBAD *mom* was introduced into the knock-out strains and *in vivo* activity of Mom was tested as previously described in figure 1. Plasmid DNA was isolated from vector control cells lacking mom expression and used as positive control for digestion with HgaI. **(D) Toxicity phenotype associated with *mom* overexpression.** Mom wild type and the inactive R101A mutant were cloned downstream of the glucose repressible and arabinose-inducible $P_{BAD}$ promoter in the plasmid pBAD24. Transformants were plated on LB media containing indicated concentrations of glucose or arabinose. Overexpression of wild-type Mom, but not the inactive mutant is cytotoxic to *E. coli* BW25113. **(E) Schematic of the genetic screen designed to investigate host genes involved in Mom modification.** *E. coli* BW25113 harbouring the plasmid pBAD *mom* cannot form colonies on 1% arabinose media. In contrast, cells harbouring a vector control or inactive Mom mutant plasmid form colonies. The lethality-based phenotype was exploited to set up a screen for host genes involved in Mom modification. Mom was expressed in a transposon mutagenesis library of *E. coli* BW25113 to screen for survivors resulting from mutations in genes involved in the Mom pathway. Chromosomal DNA is represented as a grey mesh and *mom* expression plasmid as blue circle. The yellow dots on the DNA represent DNA modification and not the Mom protein *per se*.

**Karambelkar et al., Supplementary Figure S5**

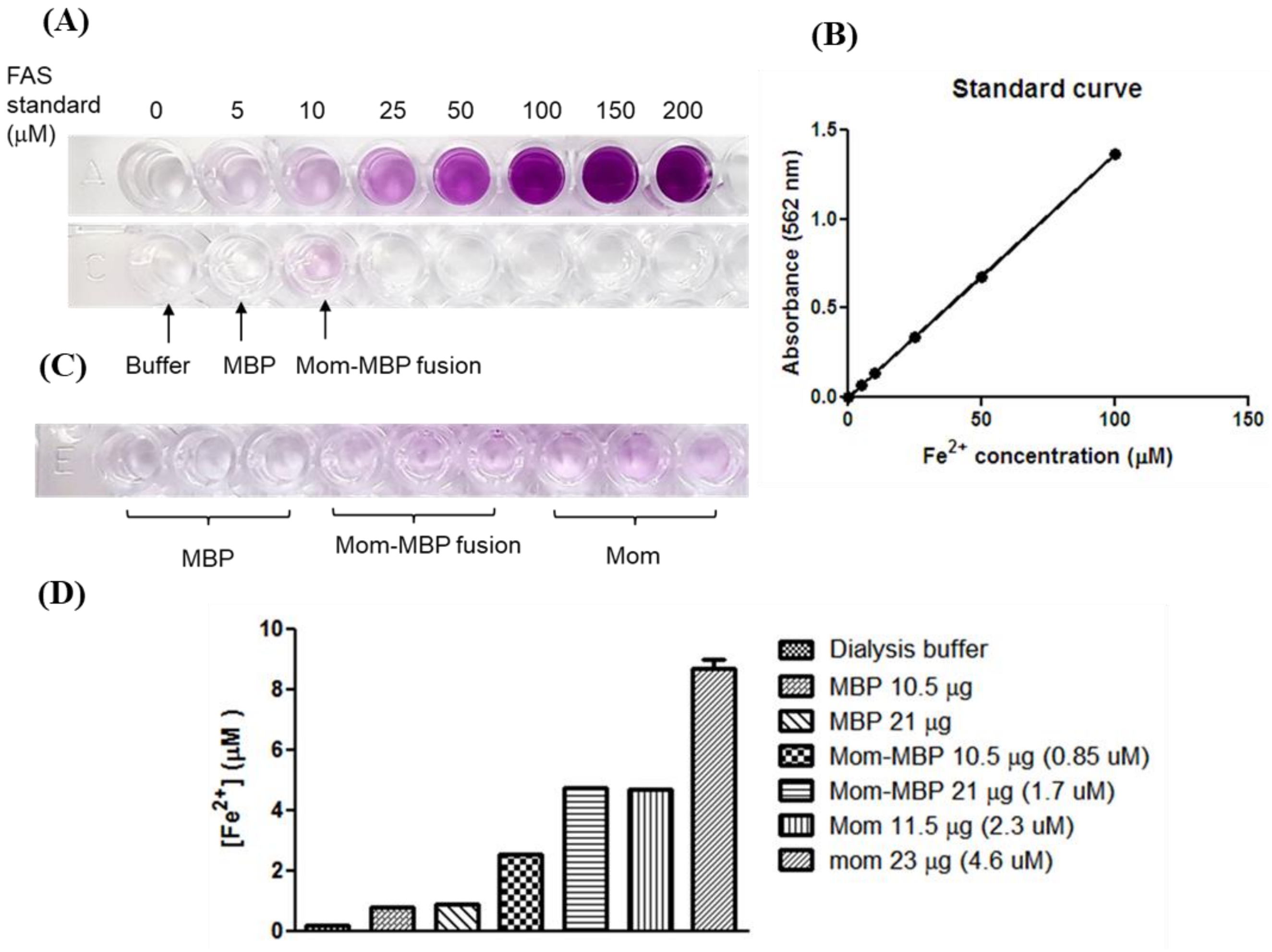


**Supplementary Figure S5. Ferrozine assay to detect binding of iron to Mom. (A)** 70 μg each of purified MBP and Mom-MBP fusion protein were subjected to ferrozine assay. Mom-MBP fusion showed a pink color indicative of intrinsically bound iron. Ferrous ammonium sulphate (FAS) standards were used to plot a standard curve within the linear range of the ferrozine assay **(B). (C)** MBP, Mom-MBP fusion and Mom were dialyzed against 10 μM FAS. Free iron was removed by dialysis and proteins were subjected to ferrozine assay. **(D)** Quantification of iron in various proteins after dialysis. Mom and Mom-MBP fusion show significantly higher levels of iron as compared to MBP.

**Karambelkar et al., Supplementary Figure S6**

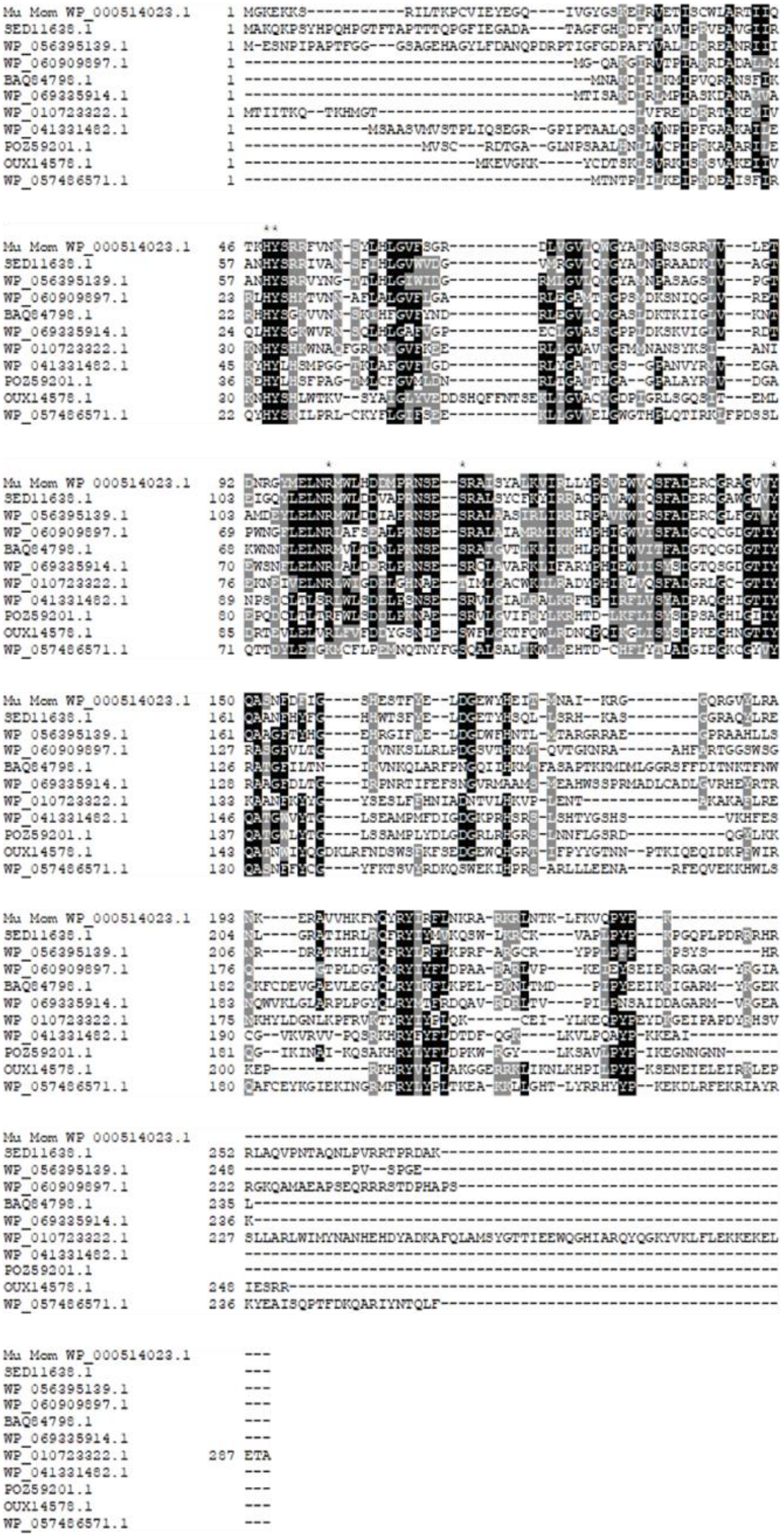

**Supplementary Figure S6. Multiple sequence alignment of Mom homologs.** Conserved residues in Mom, indicated by asterisks, were chosen for site-directed mutagenesis and resultant effects on the function of Mom were examined using plasmid restriction assays. Representative homologs of Mom obtained using an NCBI-BLASTp search (Altschul, Gish et al. 1990) were aligned using Constraint-based Multiple Alignment Tool from NCBI and the alignment was visualized using Boxshade.

**Karambelkar et al., Supplementary Figure S7**

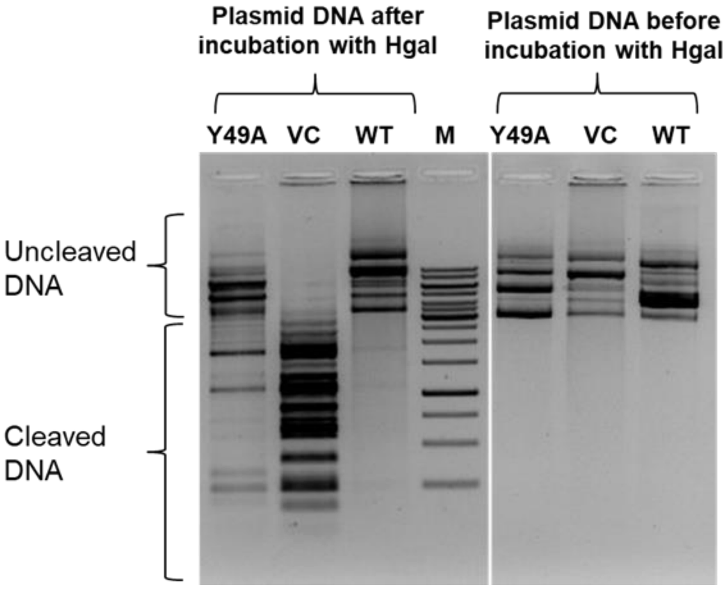


**Supplementary Figure S7. Analysis of *in vivo* activity of Y49A Mom mutant.** Plasmid DNA was isolated from *E. coli* C41 cells expressing wild type Mom or Y49A Mom as indicated. VC denotes vector control. 1 μg DNA was digested with the restriction endonuclease HgaI and products were analyzed on a 1% agarose ethidium bromide gel. Plasmid DNA purified from cells expressing Mom mutant Y49A is partially sensitive to HgaI, indicating that the mutant is compromised for DNA modification activity.

**Supplementary Table S1. Kinetic parameters for binding of Mom with DNA as determined by SPR.**

| **Ligand** | **Analyte** | **$k_{on}$ ($M^{-1}$ $s^{-1}$)** | **$k_{off}$ ($s^{-1}$)** | **$K_D$±S.D. (nM)** |
|---|---|---|---|---|
| DNA | Mom | $(3.22 \pm 0.73) \times 10^{6}$ | $(7.68 \pm 3.27) \times 10^{-4}$ | 0.234 ± 0.088 |

**Supplementary Table S2. Effect of acetyl CoA and other ligands on the thermostability of Mom.**

| **Sample** | $T_m$ °C (Mean ± SD) | |
|---|---|---|
| | **a** | **b** |
| Mom | 42.6 ± 0°C | 40.65 ± 0.49°C |
| Mom + 0.1 mM acCoA | 38.9 ± 0.14°C | 40.3 ± 0.14°C |
| Mom + 0.25 mM acCoA | 35.3 ± 0.71°C | 39.4 ± 0.14°C |
| Mom + 0.5 mM acCoA | 33.45 ± 1.2°C | 37.85 ± 0.35°C |
| Mom + 1 mM acCoA | 30.1 ± 0.14°C | 35.65 ± 0.49°C |
| Mom + 2 mM acCoA | 29.35 ± 0.92°C | 32.75 ± 0.35°C |
| Mom + 2 mM SAM | 41.95 ± 0.07°C | 40.55 ± 0.35°C |
| Mom + 1 mM CoASH | 32.9°C | |
| Mom + 1 mM malonyl CoA | 33.6°C | |
| Mom + 1 mM propionyl CoA | 43.0°C | |
| Mom + 1 mM succinyl CoA | 44.5°C | |
| Mom + 1 mM glycine | 43.0°C | |
| Mom + 1 mM NAD | 40.8°C | |
| Mom + 1 mM SAM | 42.6°C | |
| Mom + 1 mM ATP | N.D | |

[a,b] Columns a and b represent two independent experiments carried out with different batches of purified Mom stored in buffers containing 300 mM NaCl (a) and 100 mM NaCl (b), respectively. While ionic strength does influence absolute $T_m$ values, it does not affect the overall response or interaction of Mom with ligands. N.D indicates no detectable binding.

**Supplementary Table S3. Single gene knock-out strains of rSAM genes used in this study.** *E. coli* BW25113 and all the knock-out strains were procured from the Coli Genetic Stock Center, Yale University. All the knock-outs have been generated in the BW25113 (*rrnB3* Δ*lacZ4787 hsdR514* Δ(*araBAD*)*567* Δ(*rhaBAD*)*568 rph-1*) background and carry kanamycin resistance marker.

| Strain name | Description | Strain name | Description | Strain name | Description | Strain name | Description |
|---|---|---|---|---|---|---|---|
| BW25113 | wild type | JW2690-1 | Δ*hycF* | JW1471-1 | Δ*fdnH* | JW3545-1 | Δ*ysaA* |
| JW2466-1 | Δ*hyfA* | JW5501-1 | Δ*ygiQ* | JW2922-1 | Δ*yggW* | JW1620-1 | Δ*rsxB* |
| JW2192-1 | Δ*napH* | JW0878-1 | Δ*dmsB* | JW0758-1 | Δ*bioB* | JW1462-3 | Δ*narY* |
| JW3953-2 | Δ*thiH* | JW5468-1 | Δ*ygfS* | JW0764-2 | Δ*moaA* | JW5648-1 | Δ*yiaY* |
| JW2029-1 | Δ*wcaL* | JW3838-1 | Δ*hemN* | JW3864-1 | Δ*fdoH* | JW2473-1 | Δ*hyfH* |
| JW1384-1 | Δ*paaB* | JW0317-1 | Δ*yahK* | JW2437-2 | Δ*eutG* | JW3958-1 | Δ*thiC* |
| JW3486-1 | Δ*yhjA* | JW0623-1 | Δ*lipA* | JW2964-1 | Δ*hybA* | JW4033-1 | Δ*nrfC* |
| JW5248-1 | Δ*marR* | JW0885-1 | Δ*pflA* | JW1664-1 | Δ*ydhY* | JW3591-4 | Δ*Tdh* |
| JW2546-1 | Δ*yfhL* | JW3924-1 | Δ*pflC* | JW4106-3 | Δ*yjeK* | JW1492-1 | Δ*ydeM* |
| JW1228-1 | Δ*adhE* | JW0807-1 | Δ*ybiW* | JW2452-2 | Δ*aegA* | JW2501-1 | Δ*rlmN* |
| JW2748-1 | Δ*queE* | JW2836-1 | Δ*xdhC* | JW1216-2 | Δ*narH* | JW4196-3 | Δ*nrdG* |
| JW2683-1 | Δ*hydN* | JW5469-1 | Δ*ygfT* | JW4204-1 | Δ*pyrB* | JW1581-1 | Δ*ynfG* |
| JW0808-2 | Δ*ybiY* | JW5271-1 | Δ*ydhX* | JW2770-1 | Δ*fucO* | JW2134-1 | Δ*preA* |
| JW2276-1 | Δ*nuoI* | JW0819-1 | Δ*rimO* | JW3650-1 | Δ*yidF* | JW2563-1 | Δ*grcA* |
| JW2694-2 | Δ*hycB* | JW0658-1 | Δ*miaB* | JW2863-2 | Δ*fldB* | JW4342-1 | Δ*yjjW* |
| JW2193-2 | Δ*napG* | JW3178-1 | Δ*yhcC* | JW5522-1 | Δ*tdcE* | JW0977-2 | Δ*yccM* |
| JW2649-1 | Δ*nrdI* | JW2237-2 | Δ*glpC* | JW5594-1 | Δ*aslB* | JW5276-1 | Δ*ydiQ* |

**Supplementary Table S4. Bacterial strains used in this study.**

| **Strains** | **Characteristics** | **Reference** |
|---|---|---|
| *E. coli* MG1655 | $F^{-}$ $\lambda^{-}$ *ilvG*$^{-}$ *rfb*$^{-}$ 50 *rph*-1 | laboratory stock |
| BW25113 | *rrnB3* Δ*lacZ4787 hsdR514* Δ(*araBAD*)*567* Δ(*rhaBAD)568 rph-1* | *E. coli* Genetic Stock Center |
| BL21(DE3) | $F^{-}$ *ompT hsdS*$_B$($r_B^{-}$ $m_B^{-}$) *gal dcm* (DE3) | laboratory stock |
| DH10B | Δ(*mrr-hsd rms-mcrBC) mcrA recA1* | laboratory stock |
| TG1 | Δ(*hsdMS-mcrB*)5 Δ(*lac-proAB*) *supE thi-1* F'[*lacI*$^q$ *lacZ*ΔM15 *proAB*$^+$ *traD36*] | Kind gift from Prof. Umesh Varshney |
| LE392 | *hsd*R514($r_k^{-}$, $m_k^{+}$), *gln*V(*sup*E44), *try*T (*sup*F58), *lac*Y1 or Δ(*lac*IZY)6, *gal*K2, *gal*T22, *met*B1, *trp*R55 | Kind gift from Prof. Umesh Varshney |
| JW1859-1 | *rrnB3* Δ*lacZ4787 hsdR514* Δ(*araBAD*)*567* Δ(*rhaBAD)568 rph-1* Δ*cmoA* | *E. coli* Genetic Stock Center |
| JW1860-1 | *rrnB3* Δ*lacZ4787 hsdR514* Δ(*araBAD*)*567* Δ(*rhaBAD)568 rph-1* Δ*cmoB* | *E. coli* Genetic Stock Center |
| JW3975-3 | *rrnB3* Δ*lacZ4787 hsdR514* Δ(*araBAD*)*567* Δ(*rhaBAD)568 rph-1* Δ*aceA* | Kind gift from Prof. J. Gowrishankar |
| JW3974-1 | *rrnB3* Δ*lacZ4787 hsdR514* Δ(*araBAD*)*567* Δ(*rhaBAD)568 rph-1* Δ*aceB* | Kind gift from Prof. J. Gowrishankar |

**Supplementary Table S5. Sequences of primers used in this study.**

| Primer | Sequence 5'-3' |
|---|---|
| GS4 Forward | CCACGGATCCGTAATACAGATCG |
| MOM CTDEL REV | GGCCACGGATCCTCAATGATGATGATGATGATGTATCTCGTGATACC A |
| VNC2 | ATCCCGGCCCCGGACCTGGAATC |
| STS top | TATTGGGCGCTCTTCCGCTTCCTCGCTCACTG |
| STS bottom | ATAACCCGCGAGAAGGCGAAGGAGCGAGTGAC |
| MOM HISREV-1 | ATGATGATGATGATGCTTAGGGTATGGCTGAACCTTG |
| MOM HISREV-2 | GGCCACGGATCCTCAATGATGATGATGATGATGCTTAGGGTAT |
| H48AMOMF | AATTATTCAGACAAAGGCCTATTCCCGCCGTTTT |
| H48AMOMI | GTGCGGGCCAGCCAGCA |
| R101AMOMF | CTATATGGAGTTGAACGCCATGTGGCTACACGAC |
| R101AMOMI | CCCCGGTTATCCGTTTC |
| Y149FMOMF | CGCGCAGGCGTTGTGTTTCAGGCGTCGAATTTT |
| Y149FMOMI | TCCGCAGCGTTCATCTG |
| S114AMOMF | GCCCCGCAACTCTGAAGCACGGGCCATCAGCTA |
| S114AMOMI | ATGTCGTCGTGTAGCCA |
| S136AMOMF | AGTGGAGTGGGTTCAGGCCTTTGCAGATGAACG |
| S136AMOMI | GACGGATACAGTAATCT |
| D139AMOMF | GTTCAGTCCTTTGCAGCTGAACGCTGCGGACGC |
| D139AMOMI | CCACTCCACTGACGGAT |
| L69AMOMF | ATTCAGCGGACGCGATGCGGTTGGCGTTCTCCAG |
| L69AMOMI | ACCCCCAGGTGAAGGTA |
| C142AMOMF | CTTTGCAGATGAACGCGCCGGACGCGCAGGCGTT |
| C142AMOMI | GACTGAACCCACTCCAC |
| Y49AMOMF | TATTCAGACAAAGCACGCTTCCCGCCGTTTTGTGA |
| Y49AMOMI | ATTGTGCGGGCCAGCCA |
| PBADmomopfwd | AATTCACATATGGTAATACAGATCGATTATGCCCCA |
| pBADmomgFwd | AATTCACATATGGGAAAAGAGAAAAAATCACGCATTC |
| pBADmomrev | ACAGCCAAGCTTTCACTTAGGGTATGGCTGAACC |
| PBADmomhisrev | AGCCAAGCTTTCAATGATGATGATGATGATGCTTAGGGTATGGCTGA ACC |

**Supplementary Table S6. Gradient used for nucleoside analysis using LC/MS.**

| Retention (min) | % A (5 mM ammonium acetate pH 6.0) | % B (40% acetonitrile) |
|---|---|---|
| - | 100 | 0 |
| 0 | 100 | 0 |
| 3 | 100 | 0 |
| 4.4 | 99.8 | 0.2 |
| 5.8 | 99.2 | 0.8 |
| 7.2 | 98.2 | 1.8 |
| 8.6 | 96.8 | 3.2 |
| 10 | 95 | 5 |
| 25 | 75 | 25 |
| 40 | 50 | 50 |
| 44 | 25 | 75 |
| 47 | 25 | 75 |
| 55 | 0 | 100 |
| 58 | 0 | 100 |
| 63 | 100 | 0 |
| 80 | 100 | 0 |
